# Single-molecule localization microscopy reveals the molecular organization of endogenous membrane receptors

**DOI:** 10.1101/2025.08.18.670816

**Authors:** Patrick Eiring, Maximilian J. Steinhardt, Nele Bauer, Cornelia Vogt, Umair Munawar, Seungbin Han, Thomas Nerreter, Hermann Einsele, K. Martin Kortüm, Sören Doose, Markus Sauer

## Abstract

Super-resolution microscopy in combination with genetic labeling methods allows imaging of single proteins in cells. However, visualizing endogenous proteins on primary cells remains challenging due to the use of sterically demanding antibodies for labeling. Here, we demonstrate how immunolabeling conditions and antibody crosslinking influence the quantification and identification of membrane receptor stoichiometry on cells using single-molecule localization microscopy. We developed an optimized immunolabeling and analysis protocol and demonstrate the performance of the approach by resolving the molecular organization of endogenous CD45, CD69, and CD38 on Jurkat T cells. To demonstrate the usefulness of the method for immunotherapy applications we investigated the interaction of primary multiple myeloma cells with the therapeutic monoclonal antibodies (mAbs) daratumumab and isatuximab, and a polyclonal anti-CD38 antibody. Our approach might lay the foundation for improved personalized diagnostics and treatment with therapeutic antibodies.

**One-Sentence Summary:** Single-molecule localization microcopy quantifies the expression and resolves the stoichiometry of endogenous membrane receptors

## Main Text

In multicellular organisms, cells use a host of molecules for intra- and extracellular communication. A key component of these complex communication pathways are receptors, proteins that sense specific inputs and transmit the relevant biological information via cellular signal transduction and effector pathways, which influence nearly all biochemical or physiological functions of our body (*1,2*). Perturbation of receptor function often results in metabolic defects or incorrect signal transduction, leading to disease states making receptors prime targets for therapy (*3*). Considering the advances in target identification, screening technologies, and target validation, receptor molecules will remain one of the most promising targets for the development of specific drugs in the coming decades (*4*). Paritcualrily, membrane receptors play pivotal roles for diagnostics of tumors and personalized immunotherapies (*5,6*).

In recent years, super-resolution microscopy has been established as a powerful method for subdiffraction-resolution fluorescence imaging of cells and tissue (7). In the field of receptor research, single-molecule localization microscopy methods have been widely used to provide maps of plasma membrane constituents or associated proteins in unprecedented detail (*8–10*). Even though labeling of membrane proteins with primary antibodies is straightforward and is used day-to-day in clinical routine immunofluorescence and flow cytometry experiments, the labeling efficiency of membrane proteins by antibodies is limited by epitope accessibility and sterical hindrance. Furthermore, antibodies can concatenate membrane proteins resulting in the appearance of artificial membrane-protein nanoclusters (*11,12*). Therefore, most protein quantification studies have been performed using genetically modified cells to enable efficient labeling with fluorescent proteins or chemical tags (*13–16*). However, to improve receptor-directed diagnostics and guide tailored therapeutic treatment particularly for personalized therapies, information about the expression level and organization of endogenous membrane receptors is needed. Even though genomic and transcriptomic technologies can provide valuable information about protein expression, ultimately, the number of proteins and their distribution must be directly measured and quantified in the plasma membrane to deduce information about signaling functions, cellular communication and interactions (*17*).

In the last years, *direct* stochastic optical reconstruction microscopy (*d*STORM) (*18*) has been often used to visualize the distribution of plasma membrane receptors by immunolabeling using different labeling protocols (*9,19–21*). Here, we introduce a broadly applicable approach for quantifying the expression of endogenous membrane receptors on cells. In addition, by exploiting the localization statistics of antibodies in *d*STORM experiments our method provides information about the oligomeric state of endogenous receptors in the cell membrane. First, we systematically investigate the impact of fixation conditions, immunostaining with monoclonal and polyclonal antibodies, and how the fluorophore used for antibody labeling affects expression quantification of CD45, CD69, and CD38 on Jurkat T cells. CD45 is a receptor-like transmembrane protein tyrosine phosphatase and highly expressed on all nucleated hematopoietic cells where it is required for signal transduction (*22*). CD69 is a phosphorylated disulfide-linked homodimer expressed on the surface of human T-cells in the early steps of activation (*23*) and CD38 is a transmembrane glycoprotein with ectoenzymatic activity that is highly expressed on multiple myeloma cells (MM), a specific form of bone marrow cancer (*24*). Second, we employed the optimized protocol to quantify CD38 expression on OPM-2 and primary MM cells of patients diagnosed with multiple myeloma. We performed a comparative analysis of detectable CD38 on myeloma cells using two anti-CD38 therapeutic monoclonal antibodies (mAbs) currently employed in the treatment of MM patients, as well as a polyclonal anti-CD38 antibody.

### Optimizing the protocol for quantification of endogenous membrane proteins

Since fixatives have advantages and disadvantages concerning fixation speed, loss of epitopes and epitope unfolding, mislocalization of target proteins, and ultrastructure preservation (*25*), we first compared different fixation protocols with live cell immunolabeling using primary antibodies labeled with Alexa Fluor 647 (AF647) at a degree of labeling (DOL) of 2-3. *d*STORM imaging was performed by total internal reflection fluorescence (TIRF) microscopy to selectively image the basal plasma membrane at high signal-to-noise ratio. Images recorded using different fixation conditions showed mostly a homogeneous distribution of CD45 and CD69 in the plasma membrane of Jurkat T cells. However, some conditions clearly showed negative effects such as fixation-induced membrane disruption. For example, blebbing as a defined apoptosis feature was observed more often for formaldehyde (FA) and paraformaldehyde (PFA) fixed cells while ethanol-fixed and live cell labeled cells showed the highest signal-to-background ratios (Figs. 1A,B and figs. S1, S2). Cells were adhered to poly-D-lysine (PDL)-coated surfaces, which did not induce receptor cluster formation (fig. S3). To evaluate the potential formation of larger clusters, we calculated Ripley’s K-function of several regions of interest (ROI) on the plasma membrane (*26*) but did not observe significant clustering for CD45 and CD69 in live cell staining experiments (fig. S4). Based on our hypothesis that only small oligomers (e.g. homodimers in the case of CD69) may form, we used a density-based spatial clustering of applications with noise (DBSCAN) algorithm with customized localization analysis (LOCAN) (*27*) to study the spatial distribution of plasma membrane receptors and the distribution of the number of localizations per cluster that may indicate multimeric states (*28*). After selecting basal membrane regions that do not show folded membrane areas, localizations were grouped using the DBSCAN algorithm with appropriate parameters to ensure the analysis of isolated localization clusters (*29*).

**Fig. 1.**
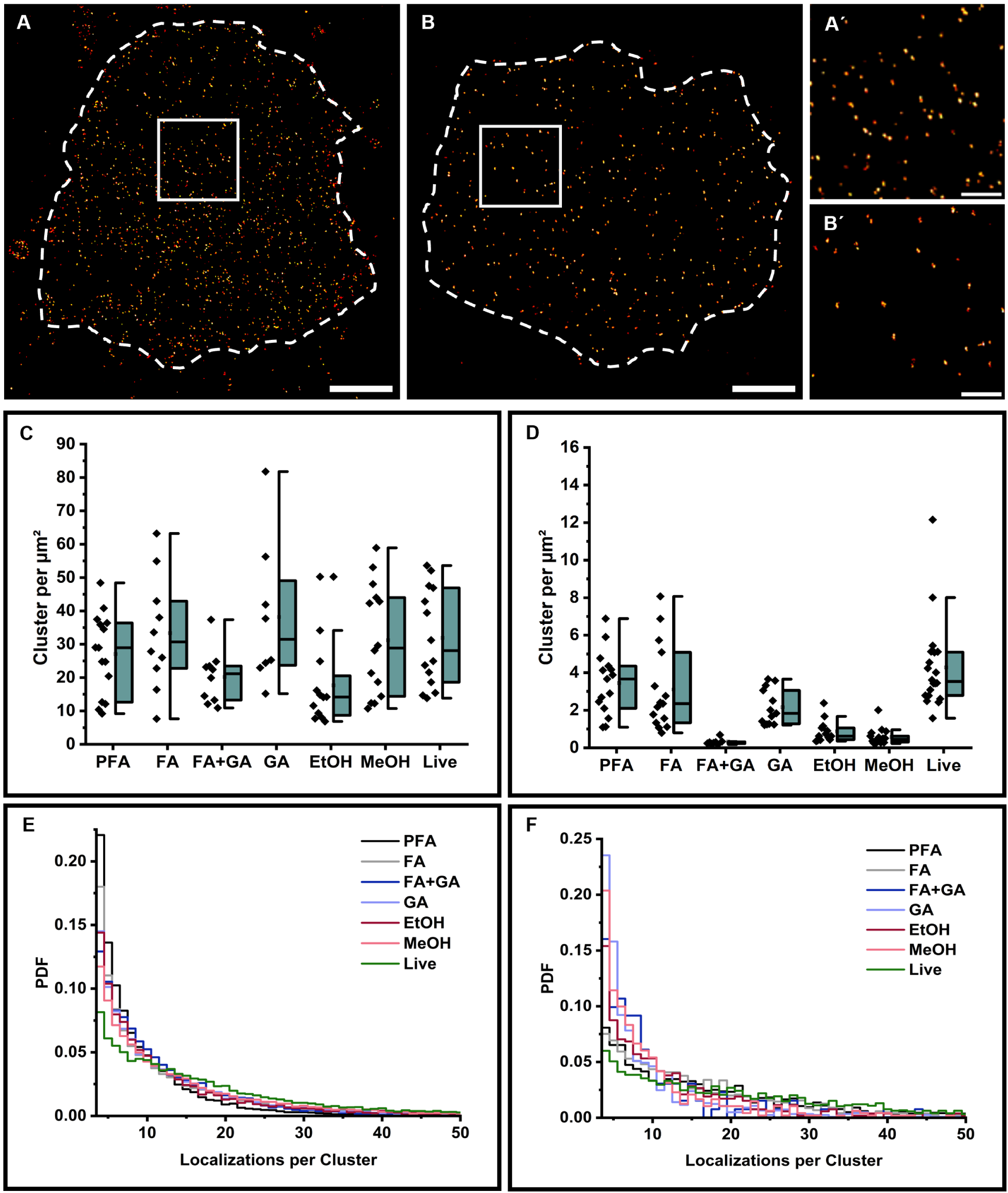
*d*STORM imaging of plasma membrane receptors. (**A,B**) *d*STORM images of CD45 labeled with the monoclonal antibody HI30-AF647 (A) and CD69 labeled with the monoclonal antibody FN50-AF647 (B) on the basal membrane of Jurkat T cells. (**Á, B.**) magnification of the area shown in A and B. (**C,D**) Corresponding localization cluster densities of CD45 (**C**) and CD69 (**D**) per µm^2^ measured for N=10-30 cells. Each localization cluster represents an isolated plasma membrane receptor. (**E,F**) Probability density functions (PDF) are shown for Jurkat T-cells stained with anti-CD45-AF647 (HI30) (**E**) and anti-CD69-AF647 (**F**). The distributions closely resemble monomeric receptor distributions. Detection of a dimeric subpopulation is achieved only when live cell immunostaining or mild fixation (FA and PFA) is applied in combination with high-affinity antibodies. Scale bars: 2 µm, magnifications 500 nm.

The resulting CD45 and CD69 localization cluster density distributions demonstrate that live cell labeling on ice followed by washing and fixation with 2% FA and 0.25% glutaraldehyde (GA) (to minimize residual mobility of membrane antigens) (*30*) detects similar amounts of receptors as alternative fixation protocols (Figs. 1C,D). Since immunostaining after fixation marks also intracellular antigens and immunotherapeutic strategies address exclusively the extracellular antigen pool, we focused our further investigations on live cell labeling on ice at 4°C, which as an additional advantage reduces antibody-binding induced antigen internalization. To ensure saturation of all accessible antigen epitopes on the plasma membrane we titrated the antibody concentration in separate experiments und used antibody concentrations of 5.0 µg/mL and 2.0 µg/mL for the detection of CD45 and CD69 antigens in the following *d*STORM experiments, respectively (fig. S5).

Next, we compared an important but often-neglected parameter of quantitative *d*STORM experiments: the photoswitching reliability of two alternative *d*STORM dyes Alexa Fluor 532 (AF532) and CF568. Although many organic dyes show photoswitching in thiol buffer, only few show a blinking behavior useful for quantitative imaging (*31,32*). Our data clearly show that AF532 and CF568 detect substantially lower amounts of CD45 at comparable DOL and attest AF647 as best suited *d*STORM dye particularly for quantitative imaging (Fig. 2A). Furthermore, *d*STORM experiments with three commonly used monoclonal anti-CD45 antibodies under identical conditions revealed significant differences in the detection efficiency possibly due to different epitope accessibilities (Fig. 2B).

**Fig. 2.**
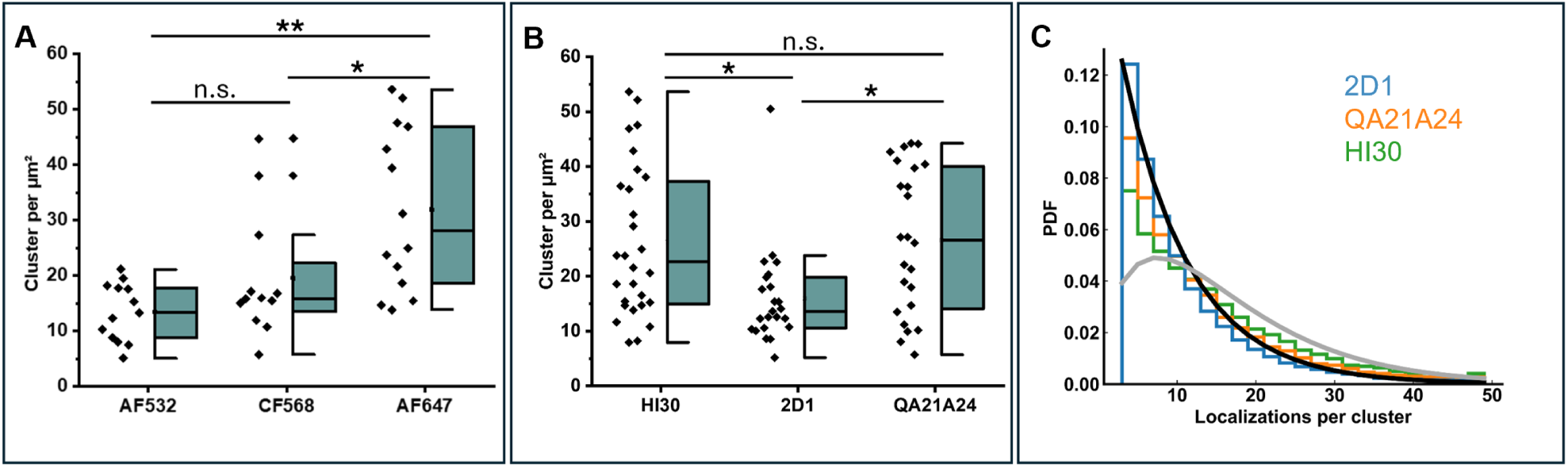
Influence of fluorophores and binding epitopes on detected cluster densities for *d*STORM data of Jurkat T cells of large passage number. **A.** Quantification of CD45 localization cluster densities using the same monoclonal antibody labeled with different dyes AF532, CF568, and AF647 measured for N=10-20 cells. We determined localization precisions of 8.4 ± 0.3 nm (mean ± std), 8.7 ± 0.2 nm, and 9.4 ± 0.5 nm for AF647, AF532, and CF568 labeled antibodies, respectively. **B.** CD45 localization cluster densities determined for three different monoclonal antibodies HI30, 2D1, and QA21A24 binding to different CD45 epitopes measured for N=20-30 cells. **C.** Histograms of the number of localizations per cluster identified in (**B**) for clone 2D1 (blue), QA21A24 (orange) and HI30 (green) are displayed as probability density function (PDF). For comparison, theoretical expectations are shown for a monomeric distribution (black) and a dimeric distribution (gray) assuming an average of 8 localizations per antibody. Histograms resemble monomeric receptor distributions for clone 2D1 and QA21A24, while for HI30 a dimeric subpopulation can be detected. The mixture distribution is only seen in cells after many passages and requires the HI30 antibody to be detected. The significance levels (**) and (*) represent p<0.01 and p<0.05, respectively. The abbreviation n.s. stands for “not significant”.

Using the optimized immunolabeling protocol with primary antibodies we tested the expression of CD38 on Jurkat T cells using the two anti-CD38 therapeutic mAbs daratumumab (DARA) and isatuximab (ISA) labeled with AF647 at a DOL of 2-3. DARA was the first CD38-targeting mAb developed and was approved for MM treatment in 2015 (*33*). ISA, which was approved for MM treatment in 2020, targets a non-overlapping epitope on the extracellular domain of CD38 (*34,35*). In addition, we used a polyclonal antibody CD38ME for CD38 quantification. *d*STORM images showed a homogenous distribution of CD38 in the plasma membrane of Jurkat T cells for all three antibodies with expression levels of 10.05 ± 1.04 (s.e.), 13.00 ± 1.64 (s.e.), and 14.60 ± 1.00 (s.e.) localization clusters per µm^2^ for DARA, ISA, and CD38ME, respectively (Figs. 3A-D). As expected, the polyclonal antibody detects the highest amount of CD38 but shows also clear signs of CD38 crosslinking, indicated by the formation of nanodomains, and a higher background signal (Fig. 3A).

**Fig. 3.**
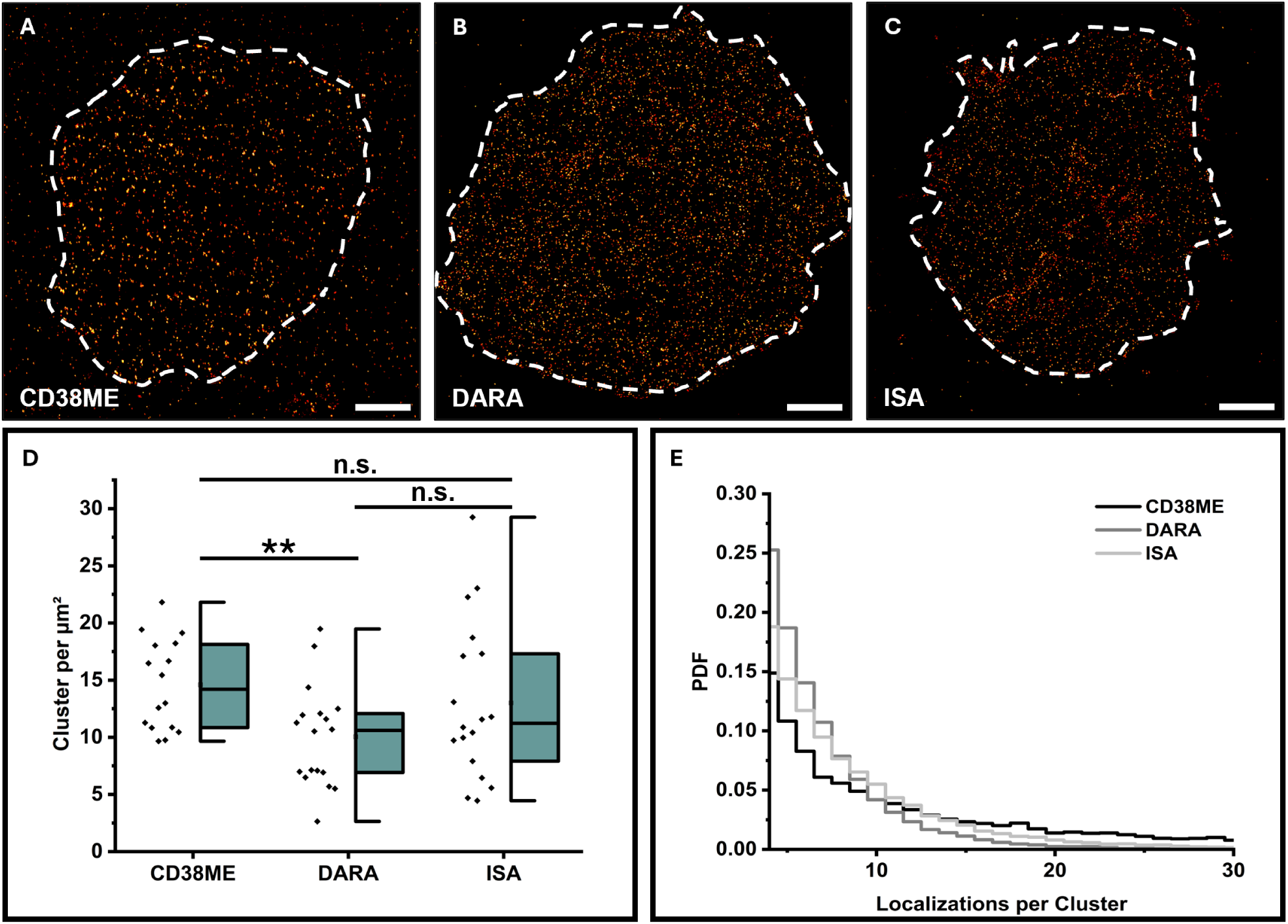
*d*STORM imaging and quantification of CD38 on Jurkat T cells. **(A-C)** Representative *d*STORM images of the basal membrane of Jurkat T cells stained with the polyclonal antibody CD38ME (**A**), DARA (**B**) and ISA (**C**). Scale bars, 2 µm. (**D)** CD38 localization clusters per µm^2^ detected for Jurkat T cells using CD38ME, DARA and ISA, all three antibodies labeled with AF647 at DOL of 2-3 (N=16-18 cells). (**E**) Probability density functions (PDFs) of the number of localizations detected per cluster for the three different antibodies. The significance level (**) represent p<0.01 and the abbreviation n.s. stands for not significant.

### Localization statistics reveal the oligomeric state of endogenous membrane receptors

Since each labeled antibody is localized multiple times in *d*STORM experiments, we investigated if the localization statistics can distinguish endogenous monomeric from dimeric plasma membrane proteins. Therefore, we analyzed the *d*STORM data and plotted the probability density function (PDF) of the number of localizations detected per spatially separated localization cluster of CD45 and CD69 measured under different immunolabeling conditions (Figs. 1E, F). Interestingly, in this set of measurements with Jurkat T cells of large passage number, the PDF determined from live cell CD45 immunolabeling experiments with the HI30 clone clearly showed a broader distribution towards higher localization numbers following the theoretical expectation for a monomer-dimer mixture (Fig. 1E). This indicates the presence of CD45 oligomers in the plasma membrane and contradicts all fixed cell immunolabeling experiments and measurements with the 2D1 or QA21A24 antibody (Fig. 2C). Indeed, different studies identified also dimeric forms of CD45 in the plasma membrane that might be important for the control of T-cell activation (*36,37*). However, while CD45 homodimers can be detected by live cell on ice HI30 immunolabeling on Jurkat T cells of large passage number (Figs. 1E, 2C), we found that CD45 always appears as pure monomer on cultured cells of small passage number (figs. S6, S7).

To investigate if the appearance of homodimers in the membrane can be induced by crosslinking of two CD45 monomers by monoclonal antibodies we performed *d*STORM experiments at varying labeling densities but did not see a change in localization cluster densities (fig. S5A) (*11,12*). Furthermore, we performed two-color *d*STORM experiments with AF647-labeled anti-CD45 primary and AF532-labeled secondary antibodies. Here, secondary polyclonal antibodies can concatenate primary antibodies bound to CD45 and induce the formation of oligomers and artificial clusters, respectively, even at 4°C (fig. S8). These data confirm that standard immunolabeling with primary and secondary antibodies cannot be used reliably for the quantification of the molecular organization of endogenous membrane proteins.

Next, we investigated the oligomeric state of CD38 on Jurkat T cells by *d*STORM using different monoclonal and polyclonal antibodies. The detected localization statistics clearly demonstrate that CD38 is detected as monomer in the plasma membrane of Jurkat T cells if labeling is performed with the therapeutic mAbs DARA and ISA (Fig. 3B-E). Staining with the polyclonal CD38ME antibody induces the formation of nanoclusters, suggesting that these structures are antibody-induced clustering artifacts (Fig. 3A,E). Finally, we investigated the expression level and oligomeric state of CD38 on OPM-2 cells (a human multiple myeloma cell line) (*38*) after labeling with a commercially available monoclonal (anti-CD38 mAb) and polyclonal antibody (anti-CD38 ME) at 4°C and 37°C, respectively and compared it to labeling with DARA. Remarkably, we did not see any signs of CD38 clustering nor indication of homodimer formation in the distribution of localization numbers per cluster using the commercially available monoclonal anti-CD38 antibody and DARA for labeling at 4°C and 37°C (figs. S9-S11). Our data gives strong evidence that labeling of live cells with monoclonal antibodies does neither induce clustering nor homodimer formation. In contrast, the polyclonal antibody CD38ME shows significant clustering, particularly when labeling is performed at 37°C (figs. S9E, S10D,E, S11).

In contrast, similar experiments performed with CD69 revealed the expected presence of homodimers especially for live cell immunolabeling but also for immunolabeling of PFA and FA fixed Jurkat T cells independent of the passage number. Under all other fixation conditions endogenous CD69 homodimers were not detected (Fig. 1F and fig. S2, S6). Next, we investigated if putative CD45 homodimers can be detected in two-color *d*STORM experiments using two different anti-CD45 antibodies. Therefore, we first immunostained living Jurkat T cells of large passage number on ice with clone 2D1 labeled with AF647, followed by staining with clone HI30 labeled with CF568. The resulting two-color images show the presence of overlapping binding events indicating that the two different clones can bind simultaneously to CD45 homodimers albeit at low efficiency (fig. S12). Furthermore, we validated that two different antibodies can bind to CD69 homodimers as indicated by the appearance of two-color binding events (fig. S13A) whereas colocalization experiments performed with anti-CD45 and anti-CD69 showed no colocalization events (fig. S13B). Unfortunately, immunolabeling of endogenous membrane receptors will not enable quantification of the monomer/homodimer equilibrium because of different epitopes addressed by the different antibodies and restricted epitope accessibilities using sterically demanding IgG antibodies. Nevertheless, our data clearly emphasize the importance of the choice of the antibody and fixation conditions used for quantification of endogenous membrane proteins.

### Impact of the antibody used for quantification on immunotherapy treatment

Recently, *d*STORM has been used to quantify CD38 with AF647 labeled DARA on 31 DARA naïve and 22 DARA resistant MM patients. In this study the mean CD38 receptor density was significantly higher in DARA-naïve patients (*39*). Exemplarily we show here the data obtained from patient 677, an IgA lambda myeloma patient, who was newly diagnosed and untreated at the time of sampling. He achieved stringent complete remission following subsequent DARA-based induction therapy. Here, *d*STORM quantified similar amounts of CD38 of 16.0 ± 1.1 (s.e.) and 19.3 ± 1.6 (s.e.) localization clusters per µm^2^ for polyclonal anti-CD38ME and DARA, respectively (Figs. 4A,B). Using the mAb ISA for labeling we detected slightly higher amounts of 22.5 ± 1.4 (s.e.) localization clusters per µm^2^ on primary myeloma cells of patient 677 (Fig. 4C,G).

**Fig. 4.**
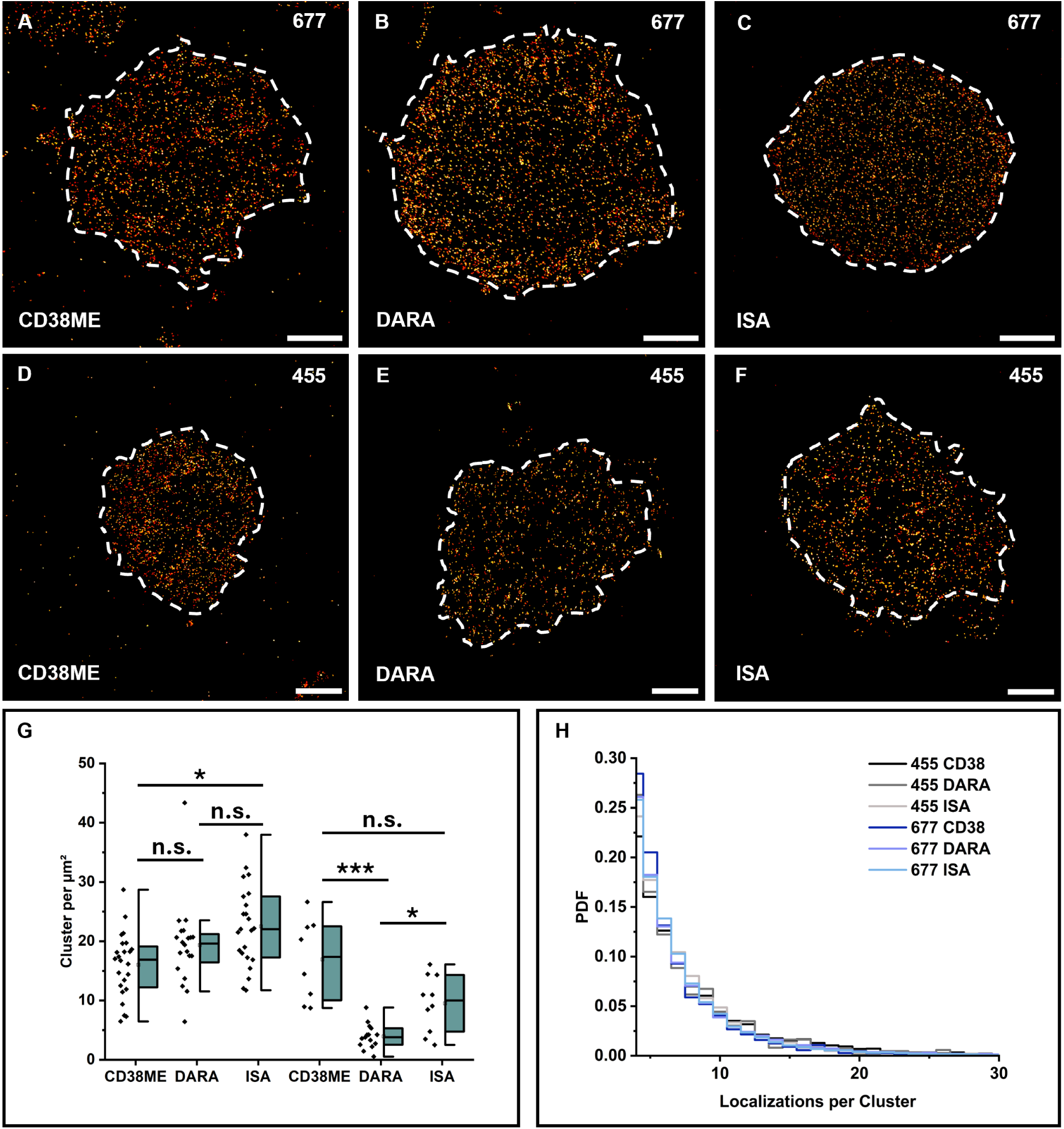
*d*STORM imaging and quantification of CD38 on patients responding to and resistant to Daratumumab treatment. **A-C.** Representative *d*STORM images of the basal membrane of Multiple Myeloma cells stained with the polyclonal antibody CD38ME (**A**), Daratumumab (**B**) and Isatuximab (**C**) are shown for patient 677, who is responding to DARA treatment. **D-F.** Representative *d*STORM images of the basal membrane of Multiple Myeloma cells stained with the polyclonal antibody CD38ME (**D**), Daratumumab (**E**) and Isatuximab (**F**) are shown for patient 455, who developed a resistance to DARA treatment. **G.** CD38 localization clusters per µm^2^ detected for patient 677 and 455 using CD38ME, Daratumumab and Isatuximab labeled with AF647 (N=8-30 cells). **H.** Probability density functions (PDFs) of the number of localizations detected per cluster for the two different patients and antibodies. Scale Bars: 2 µm. The significance levels (*) and (***) represent p<0.05 and p<0.001, respectively. The abbreviation n.s. stands for not significant.

However, MM cells from patient 455 showed DARA resistant multiple myeloma cells. He was initially well responsive to DARA but was then sampled in aggressive relapse following 12 months of continuous DARA containing therapy (*39*). Here, we identified a similar amount of CD38 with anti-CD38ME of 16.9 ± 2.5 (s.e.) localization clusters per µm^2^ but a substantially lower amount of CD38 with DARA of only 3.9 ± 0.5 (s.e.) localization clusters per µm^2^ (Figs. 4D,E). However, immunolabeling with ISA demonstrates a notably higher sensitivity in detecting CD38 molecules in this patient, identifying more than twice as many as DARA with 9.5 ± 1.5 (s.e.) localization clusters per µm^2^ (Figs. 4F,G). On the other hand, MM cells of patient 824 became resistant during DARA treatment albeit DARA is efficiently binding to MM cells (fig. S14).

Unfortunately, DARA resistance mechanisms remain poorly understood. Proposed resistance mechanisms range from intrinsic downregulation in CD38^dim^ MM cells due to genetic alterations (*39*) to prolonged blocking of CD38 epitopes by DARA (*40*). Our data show that a biallelic loss of CD38 as described recently (*38*) cannot be responsible for our observation since DARA is still binding on MM cells of patient 455 albeit with reduced binding efficiency. Our observation is particularly important as it demonstrates that the use of an alternative mAb such as ISA, provides the potential to be used advantageously for the treatment of albeit rare MM patients with acquired genetic alterations or epitope saturation that affect the DARA binding efficiency. Currently, switching CD38 antibodies after the development of resistance is not part of routine clinical practice. However, our imaging methodology may open new opportunities for at least a subset of patients, as recent studies identified point mutations with clinical impact on DARA sensitivity, such as L153H and R140G, in selected patients (*39*). Noteworthy, these mutations appear to be rare in the 1,186 patients of the CoMMpass study, and among 399 whole-genome-sequenced MM patients from our institution, only one exhibited a subclonal CD38 mutation (*41*).

## Discussion

Receptor-directed diagnostic and therapeutic strategies are increasingly used in various fields of clinical medicine, particularly in hematology and oncology. Thus, novel immunotherapeutic strategies, which allow to redirect immune cells by using artificial receptors recognizing surface antigens on tumor cells, are offering unprecedented treatment opportunities even in very advanced and relapsed refractory hematological malignancies. In addition, pharmacological inhibition of mutated and wild-type receptors and T cell redirecting therapies are increasingly used to treat solid tumors, autoimmune disorders and other inflammatory conditions (*42,43*). However, the entire field of immunotherapy relies critically on the availability of reliable expression numbers of membrane antigens. Obviously, genetic modification of cells with fluorescent proteins and chemical tags allows stoichiometric labeling and thus simplified quantification of membrane proteins (*13–16*). However, detection of endogenous tumor-associated receptors remains essential to assess target accessibility for immunotherapeutic treatment of individual patient cells. In this context, *d*STORM stands out since it allows single-molecule sensitive visualization of membrane receptors with commercially available fluorescently labeled primary antibodies. Here, it is certainly a drawback that the immunolabeling efficiency of endogenous membrane proteins with primary antibodies is unknown. Therefore, absolute expression numbers cannot be appropriately determined. However, *d*STORM imaging and analysis using the therapeutic mAb for imaging that is used for patient treatment enables quantitative measurements of expression numbers that predict immunotherapy success chances for individual patients.

Unfortunately, until now no universally accepted method for quantification of endogenous membrane receptors has been established. Therefore, we investigated the impact of sample preparation, particularly the choice of fixative, on *d*STORM quantification and the possibility to resolve the stoichiometry of endogenous membrane receptors. Our results show that fixation conditions can profoundly influence both the quantitative detection of receptors and the identification of small oligomers and thus result in false-negative results, potentially affecting clinical treatment decisions. This finding underscores the importance of standardized protocols to avoid diagnostic discrepancies, which are crucial in the context of personalized immunotherapy. While spatial distribution analysis (e.g. using Ripley’s K-function) is a widely accepted and often used method for assessing cluster formation, its utility is limited when it comes to small oligomers such as homodimers with a size below the resolution limit. However, analyzing localization data with a DBSCAN-clustering algorithm combined with customized localization analysis can tackle this challenge and deliver robust information about the oligomeric state of endogenous membrane receptors. To conclude, our approach enables important insights into the interaction of endogenous membrane proteins and antibodies, which can be used advantageously for the improvement of personalized immunotherapy treatments and the design of more effective therapeutic antibodies.

## Materials and Methods

### Cell culture of Jurkat T cells and OPM-2 cells

Human T-lymphoblast (Jurkat E6-1, Cell Lines Service GmbH, #300223) and OPM-2 (DSMZ, #ACC 50) were cultured in RPMI-1640 (Sigma-Aldrich, #R8758) containing 10% FCS, 100 U/mL penicillin and 0.1 mg/mL streptomycin at 37°C and 5% CO_2_. Cells were maintained at a maximum density of ∼2 x 10^6^ cells/mL in standard T25 culture flasks (Sarstedt # 83.3910.502).

### Antibody-dye conjugation

Daratumumab (Darzalex, Johnson & Johnson) and Isatuximab (Sarclisa, Sanofi) were kindly provided by the Pharmacy of the University Hospital of Würzburg, anti-CD38 (clone: HIT2 #303502, Biolegend), anti-CD38 ME (Cytognos), anti-CD45 antibodies (clone: 2D1 #368502; clone: HI30 #304002; clone: QA21A24 #384402, Biolegend) and anti-CD69 (clone: FN50 #310902) were self-labeled with AF647-NHS (Thermofisher, #A20006). CD45 (HI30) was additionally labeled with CF568-NHS (Merck, #SCJ4600027) for two-color experiments. The antibodies were labeled at RT for 2 h in 100 mM sodium bicarbonate (Fisher Scientific, 144-55-8, pH 8.5) following the manufacturers standard protocol. Briefly, 50 µg of antibody was reconstituted in sodium bicarbonate buffer using 0.5 mL spin desalting columns (40K MWCO, Thermofisher, #87766). To get an average DOL of 2-3 a 5× dye excess was used. All antibodies were purified and washed using additional spin desalting columns to remove unbound dye. Finally, antibody concentration was determined measuring absorption (A) at 280 nm and 568 nm or 647 nm (A_dye_) with a nanophotometer respectively and calculated according to the following formula with ε being the extinction coefficient and CF the correction factor of the dye at 280 nm.

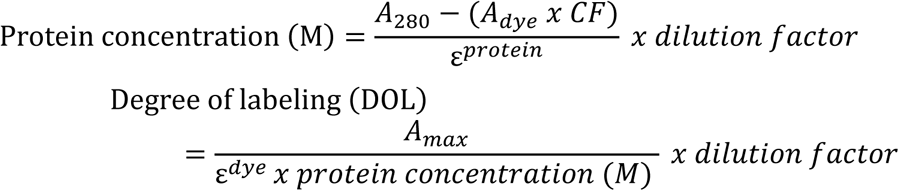

### Coating dependent cluster formation

Jurkat T cells (DSMZ, Braunschweig, Germany) modified to stably express a ZAP70-GFP fusion protein were kindly provided by Dr. Thomas Nerreter (University Hospital, Würzburg). Cells were seeded at a concentration of 2.5 × 10^5^ cells/well into eight-well chambered glass slides with a high-performance cover glass bottom (8-well Chambered Coverglass System #1.5 High Performance Cover Glass (0.17 ± 0.005 µm), Cellvis) coated with either 0.1 mg/mL poly-D-lysine (PDL) (Sigma-Aldrich, #P6407) or 5 µg/mL anti-CD3 (clone: UCHT1 #300402, Biolegend) antibody. After allowing cells to adhere for 2 hours in an incubator, the basal membrane was imaged using a Zeiss LSM 700 Confocal Microscope equipped with an oil immersion objective (PlanApochromat 63× 1.40 NA, Zeiss). Imaging was performed with a confocal pinhole diameter of 1.0 Airy unit using a 488 nm laser (10 mW) set to 5% intensity and a pixel dwell time of 5 µs to visualize ZAP70-GFP cluster formation.

### Live cell staining of Jurkat T cells and MM cells

Jurkat T cells, OPM-2 and MM cells were seeded at concentrations of 2.5 × 10^5^ cells/well into PDL coated eight-well chambered glass slides with a high-performance cover glass bottom and allowed to adhere in the incubator. After the cells successfully attached to the glass surface, they were incubated at room temperature for 5 mins before being transferred to ice. The media was replaced with ice cold PBS containing either anti-CD45 or anti-CD69 antibodies for Jurkat T cells in concentrations ranging from 0.5 µg/mL-10 µg/mL or anti-CD38 ME (Cytognos), Daratumumab or Isatuximab for MM and Jurkat T cells with a working concentration of 7.5 µg/mL for 30 mins, respectively. Additionally, a monoclonal anti-CD38 antibody was used for staining of the OPM-2 cells. After washing, cells were fixed for 15 mins with 2% methanol-free formaldehyde (FA) (ThermoFisher, #28906) and 0.2% glutaraldehyde (GA) (Sigma-Aldrich, #G5882-10×1ML), again washed with PBS and stored at 4°C in the dark or measured directly.

### Post-fixation staining of Jurkat T cells

Jurkat T cells were seeded at concentrations of 2.5 × 10^5^ cells/well into poly-D-lysine (PDL) (Sigma-Aldrich, #P6407) coated eight-well chambered glass slides with a high-performance cover glass bottom ((8-well Chambered Coverglass System #1.5 High Performance Cover Glass (0.17 ± 0.005 µm), Cellvis) and allowed to adhere in the incubator. After successful attachment cells were either fixed with 4% freshly solved methanol free PFA, 4% FA, 4% FA + 0.25% GA, 2% GA for 15 minutes at RT or with ice cold pure Methanol or Ethanol at −20°C for 20 minutes. Afterwards cells were washed thrice with PBS before being stained with previous defined saturating concentrations of 5 µg/mL anti-CD45 antibodies or 2.5 µg/mL anti-CD69 antibody for 30 mins. After washing, cells were postfixed for 15 mins with 2% methanol-free formaldehyde (FA) and 0.2% glutaraldehyde (GA), again washed with PBS and stored at 4°C in the dark or directly measured.

### *d*STORM Imaging

Imaging of membrane receptors was performed utilizing an inverted wide-field fluorescence microscope (IX-71; Olympus) equipped with a nose piece stage for improved stability. For excitation of AF647 coupled antibodies, a 640 nm diode laser (Cube 640-100C, Coherent) in combination with a clean-up filter (laser clean-up filter 640/10, Chroma) was used. The laser beam was focused onto the back focal plane of the oil-immersion objective (60×, NA 1.45; Olympus). Emission light was separated from the illumination light using a dichroic mirror (HC 560/659; Semrock) and spectrally filtered by a band-pass filter (FF01-679/41-25, Semrock). Images were recorded with an electron-multiplying CCD camera chip (iXon DU-897; Andor). Pixel size for data analysis was measured to be 128 nm. For imaging 256 × 256-pixel areas with TIRF excitation with an exposure time of 20 ms (frame rate 50 Hz) and irradiation intensity of 2.5 kW/cm^2^ were used. Images were recorded for 15k frames (5 mins) for each measurement. To induce photoswitching of AF647 a PBS based buffer (pH 7.4) containing 100 mM β-mercaptoethylamin (Sigma-Aldrich, M6500) was used.

### Preparation of TetraSpeck probe

Cellvis 8-well chambers were cleaned by incubating them in 1M KOH for 10 min, followed by washing twice with ddH_2_O. The chambers were further cleaned with absolute ethanol (100%) for 20 min, removed, and air-dried under a clean bench. To coat the wells, 150 µl of 0.1 mg/ml poly-D-lysine (PDL) solution was added to each well and incubated for 1 h. The PDL solution was then removed, and the wells were rinsed once with ddH_2_O. TetraSpeck microspheres (Invitrogen, T7279) were prepared by diluting them 1:100 in an 80 mM PIPES buffer (Sigma, P1851) containing 1 mM MgCl_2_ and 1 mM EGTA at pH 6.8. The diluted solution was sonicated for 30 minutes to ensure proper dispersion of the beads. Following sonication, 150 µl of the solution was added to each well and incubated for 1 hour. Finally, the wells were washed three times with 1x PBS to remove unbound microspheres.

### Two-color *d*STORM

Two-color *d*STORM imaging was performed on the same homebuilt widefield setup as described before. First, both camera channels were pixel precise aligned by placing the selfmade TetraSpeck sample on the microscope stage and adjusting the beam splitter to align the centers of both camera chips on the same fluorescent speck. Afterwards, the two-color CD45 sample was put on stage and after focusing on the basal layer the setup was left for a certain time until no z-dimensional drift was observed anymore. The image stacks of 15,000 frames were first acquired for anti-CD45-AF647 (2D1) and followed by anti CD45-CF568 (HI30). The sample was removed and replaced by the TetraSpeck sample, which was imaged (100 frames with 50 ms exposure time) until at least a few TetraSpecks were located in each corner and in the middle of the camera chip. This workflow was repeated after each two-color image to ensure a high precision of the later created elastic transformation by the Fiji plugin “bUnwarpJ” for both cameras. Photoswitching was achieved by using a slightly adjusted imaging buffer of 100 mM MEA in PBS pH 7.7.

### Data analysis

The recorded *d*STORM images were reconstructed with rapidSTORM 3.3. Localization data acquired in *d*STORM measurements were filtered to remove background noise with less than 800 photons for CF568 and AF647 and 600 photons for AF532. For analysis of each *d*STORM image an appropriate region of interest (ROI) at the basal membrane of the cell, was chosen. Localization analysis was carried out with customized python scripts based on Locan^23^ and the scientific python stack. We calculated and displayed Ripley’s H-function, a normalized Ripley’s K-function, using Locan as previously described^22,51^. Computation was carried out for each ROI indicating individual cells without edge correction, yielding an average H-function and 95% confidence intervals. This experimental data was compared to H-functions and their 95% confidence intervals computed from 100 simulated data sets with localizations distributed on the same ROIs, and with identical number of localizations in each ROI, according to complete spatial randomness or a Neyman-Scott process resembling *d*STORM data. The Neyman-Scott clustering process has homogeneously distributed parent events with each parent having n offspring events, where n is Poisson distributed with mean 15, and with the offspring positions having a Gaussian offset with a standard deviation of 10 nm. The maximum of the H-function indicates a distance that is between cluster radius and diameter and thus provides an estimate for the average cluster size. For cluster analysis, the DBSCAN clustering algorithm was employed to group detected localizations with ε = 20 and minPoints = 3, determined via analysis of synthetic datasets^22^. These parameters facilitated quantification of detected localization within a certain distance, providing insights into existing oligomeric states and addressable receptors on the cell surface. Measurement of the average number of localizations per localization cluster for individual antibodies (as seen in samples labeled with low antibody concentrations) yielded an expected ∼7-8 localizations per antibody under the given acquisition conditions. To determine the receptor stoichiometries that are below the resolution limit of *d*STORM, we used a probability density function (PDF) to analyze the distribution of localizations within DBSCAN-identified clusters. This analysis allowed us to estimate whether receptors are likely to be monomeric, dimeric, or part of a higher-order oligomer. Experimental data were compared to theoretical expectations^24^ as described by a negative binomial distribution with probability mass function (PMF)

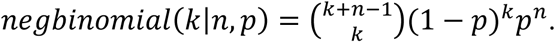

with parameters *n* ∈ (1,2) representing monomers and dimers, respectively, and *p* = 1⁄(1 + *l*) where *l* is the mean number of localizations per cluster from a monomer.

Experiments were conducted using a single coherent dataset sequentially to eliminate instrumental variations. Box plots were utilized to depict data distribution, with boxes representing values between the first and third quartiles (25% −75%) and the median indicated by a center line. Whiskers extend to 1.5 times the interquartile range. Statistical analysis was conducted with a student’s t-test using OriginPro software version 2021b, with a significance level set at p < 0.05. Reported p-values are denoted with asterisks: * for p < 0.05, ** for p < 0.01, and *** for p < 0.001.

If no p-value is presented in a specific panel, it suggests that no statistically significant differences were observed between the compared groups.

## ACKNOWLEDGEMENTS

The authors thank Elke Maier for cell culture support.

## Funding

K.M.K. received research funding from Janssen-Cilag, the Stifterverband and from the Deutsche Forschungsgemeinschaft (DFG, German Research Foundation, TRR 387 and KFO 50001). S.D. received funding from the Deutsche Forschungsgemeinschaft (DFG, German Research Foundation) project DO1257/4-1. T.N., P.E. and M.S. received funding from SFB - TRR 338/1 2021 – 452881907 and the German Ministry for Science and Education (BMBF, Bundesministerium für Bildung und Forschung, Grant #13N15986). K.M.K. received funding from the Deutsche Krebshilfe via MSNZ.

### Author contributions

P.E., S.D. and M.S. conceived and designed the project. S.D. and M.S. supervised the project. C.V. prepared MM samples. P.E. and N.B: prepared Jurkat and OPM-2 samples. P.E. performed all *d*STORM experiments. K.M.K., H.E., T.N., U.M., S.H. and M.J.S supported all MM experiments. P.E. and S.D. performed data analysis. P.E., S.D., K.M.K and M.S. wrote the manuscript. All authors revised the final manuscript.

## Competing interests

The authors declare no conflict of interest.

### Data and Materials availability

The data that support the findings of this study will be provided by the corresponding author upon reasonable request.

## SUPPLEMENTARY MATERIALS

Figures S1 to S14

Supplementary Materials for

**Fig. S1.**
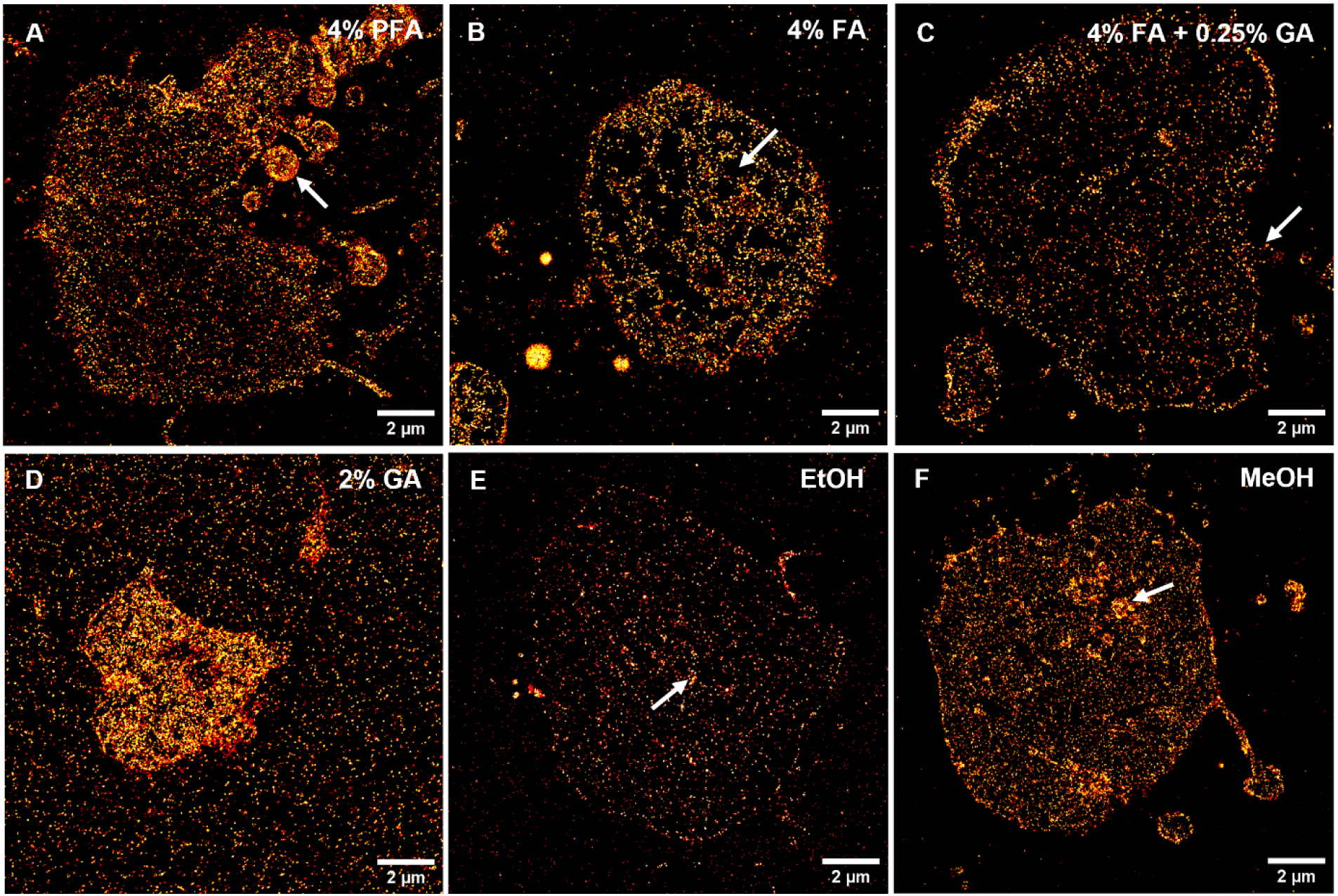
Representative *d*STORM images of the basal membrane of Jurkat T cells prefixed with various fixatives following staining with CD45-AF647 (HI30). Fixation with 4% paraformaldehyde **A.** and 4% formaldehyde (**B**) leads to more pronounced apoptotic features, such as visible membrane blebbing or membrane disruptions (white arrows). **C.** A combination of 4% formaldehyde with 0.25% glutaraldehyde on the other side shows a high signal-to-noise ratio with only minor disruptions. **D.** Fixation with 2% glutaraldehyde induces high unspecific binding of the antibody to the glass surface. Cells fixed with 100% ethanol (**E**) and 100% methanol (**F**) exhibit a high signal-to-noise ratio, however, ethanol-fixed cells show reduced binding and intracellular staining. Arrows indicate the described features in each respective image.

**Fig. S2.**
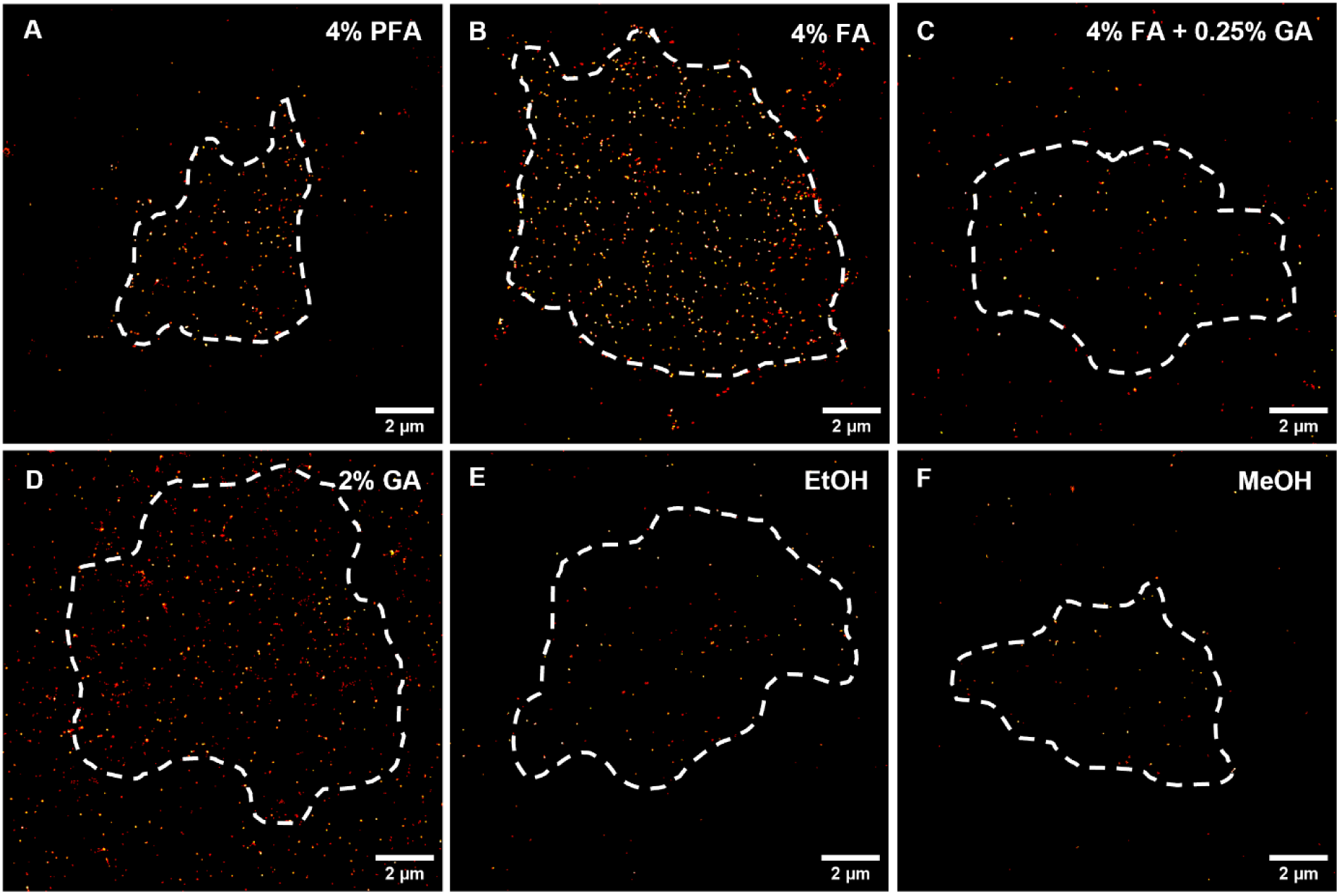
Representative *d*STORM images of the basal membrane of Jurkat T cells prefixed with various fixatives following staining with CD69-AF647. Clear detection of CD69 molecules was observed with 4% paraformaldehyde (**A**) and 4% formaldehyde (**B**). However, protocols using 4% formaldehyde + 0.25% glutaraldehyde (**C**), 100% ethanol (**E**), and 100% methanol **(F**) resulted in failed or minor detection of CD69 expression. Additionally, a significantly higher amount of nonspecifically bound CD69 antibodies to the glass surface was observed with 2% glutaraldehyde prefixation (**D**).

**Fig. S3.**
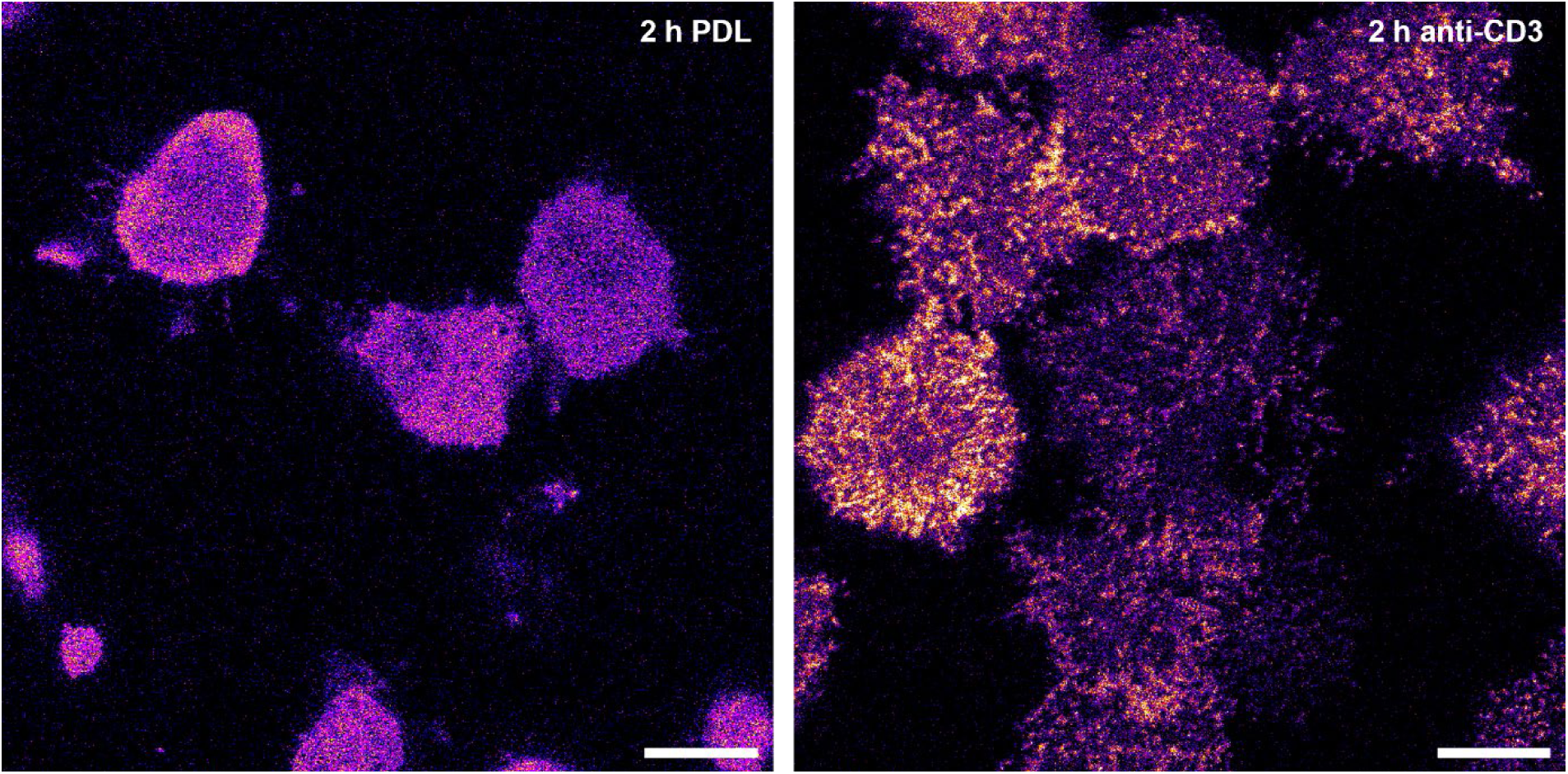
Confocal laser scanning images of Jurkat T ZAP70-GFP cells after 2 h on a PDL coated surface (left) and an anti-CD3-coated surface (right). Intracellular ZAP70-GFP remains homogeneously distributed on PDL, while binding to anti-CD3 antibodies activates the Jurkat T cells as indicated by ZAP70-GFP nanocluster formation.

**Fig. S4.**
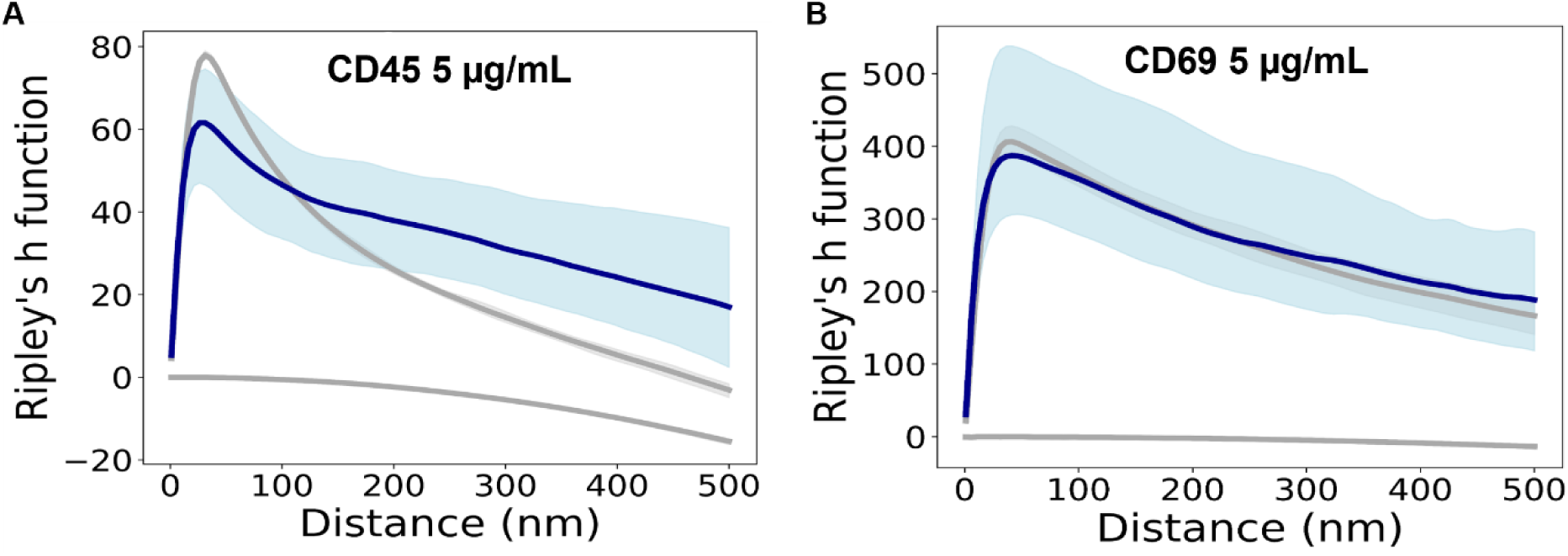
Ripley’s H-function analysis for *d*STORM data of Jurkat T cells of low passage number immunostained for CD45 and CD69. No significant clustering was observed for anti-CD45 (HI30) (**A**) and anti-CD69 (**B**) antibody staining on Jurkat T cells. Furthermore, no difference in cluster size between the two receptors was detected. For comparison with experimental data from 8 and 6 cells (blue), Ripleýs H function for simulated data with spatial distributions following complete spatial randomness (lower grey lines) or a clustered Neyman-Scott process (upper grey lines) in identical ROIs are displayed with 95% confidence intervals (light blue and gray regions; if not shown confidence intervals are smaller than the line width).

**Fig. S5.**
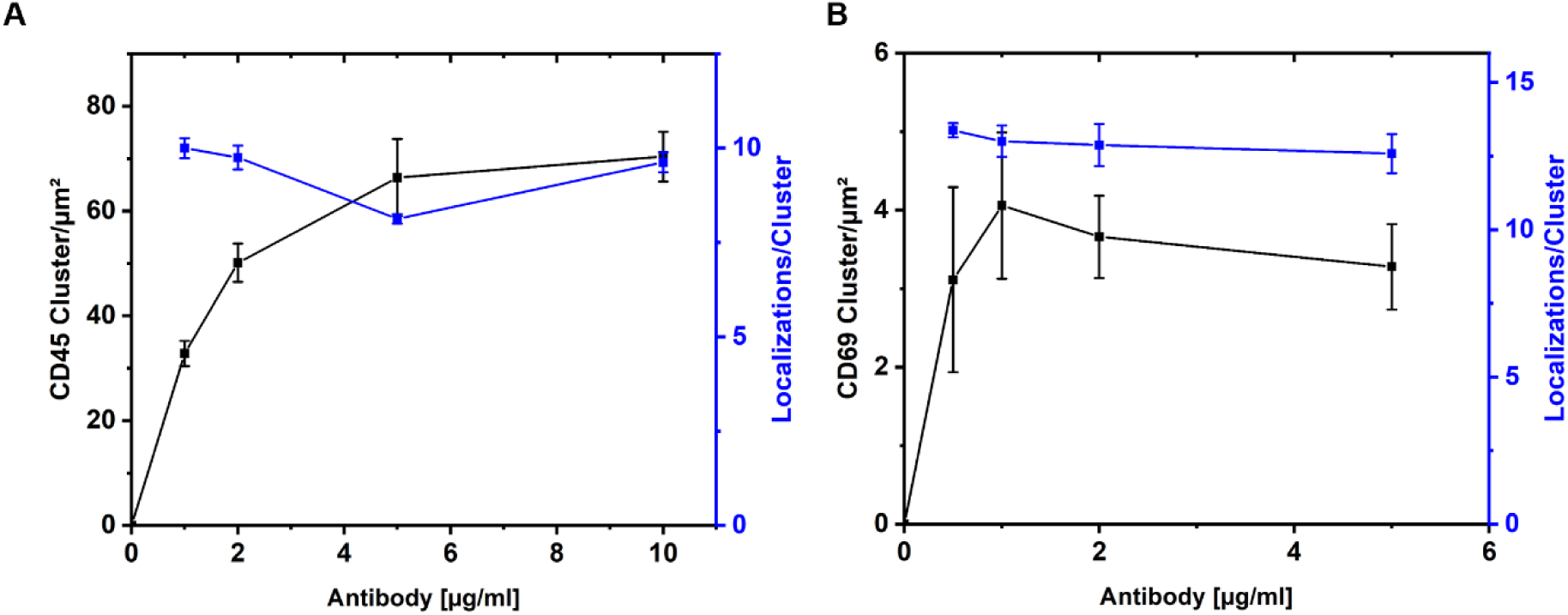
Average number of CD45 and CD69 localization clusters detected on the plasma membrane of Jurkat T cells at varying antibody concentrations. **A,B,** The data indicate that an antibody concentration of 5 μg/mL and 2 µg/mL is sufficient to label all accessible CD45 (**A**) and CD69 (**B**) epitopes on the plasma membrane, respectively. Furthermore, the number of localizations detected per cluster remains consistent across varying concentrations of the antibodies, supporting the conclusion that the monoclonal CD45 (**A**) and CD69 (**B**) antibody does not induce artificial clustering.

**Fig. S6.**
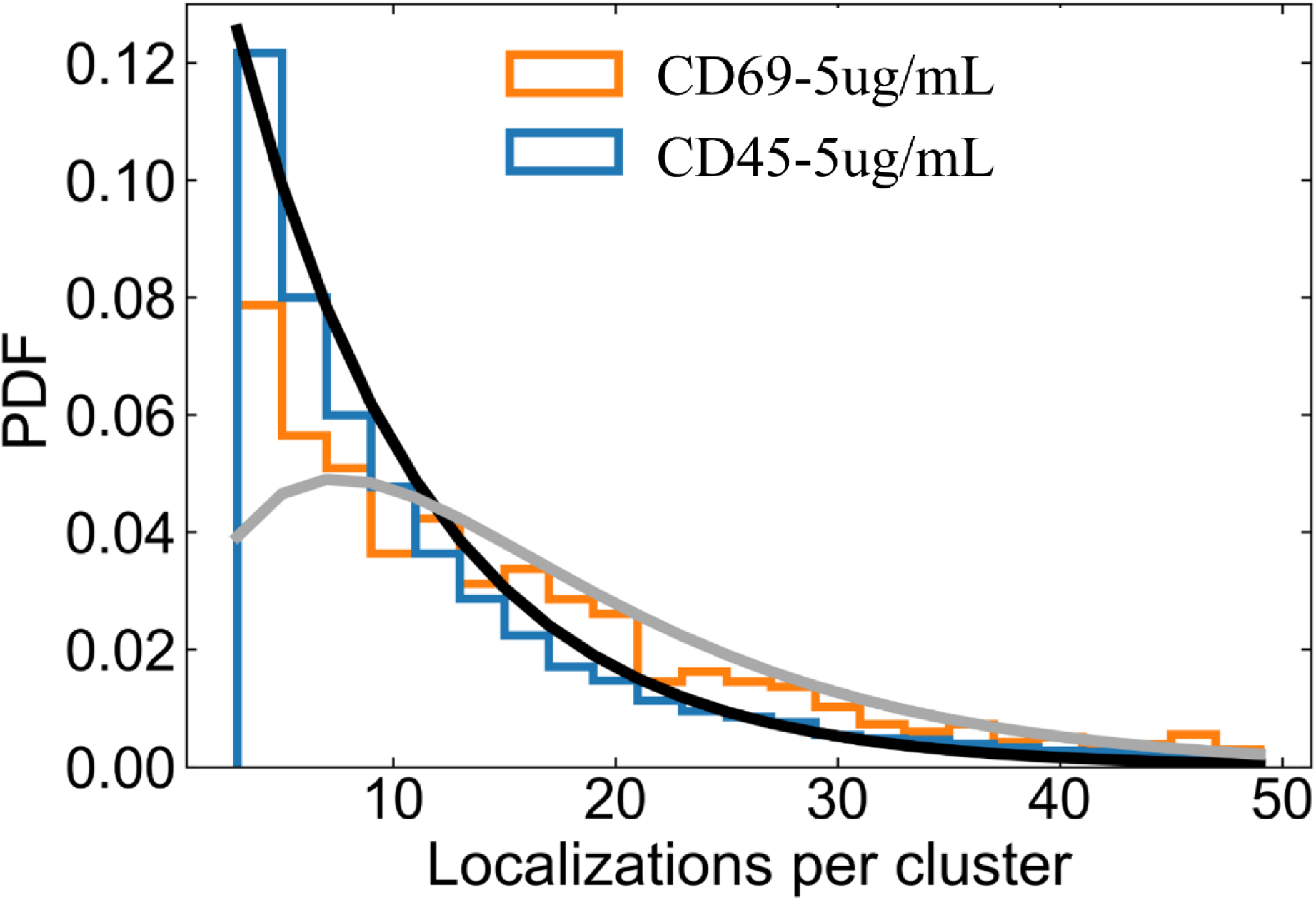
Probability distribution for the number of localizations per cluster for *d*STORM data of Jurkat T cells of low passage number immunostained for CD45 and CD69. The same *d*STORM localization data as analyzed in fig. S4 was clustered with a DBSCAN algorithm. Histograms of the number of localizations per cluster for anti-CD45 (HI30) (blue) and anti-CD69 (orange) antibody staining on Jurkat T cells are displayed as probability density function (PDF). For comparison, theoretical expectations are shown for a monomeric distribution (black) and a dimeric distribution (gray) assuming an average of 8 localizations per antibody. For CD45 (HI30) the distribution resembles that of monomers. For CD69 a mixture distribution of monomers and a more pronounced dimeric population is observed.

**Fig. S7.**
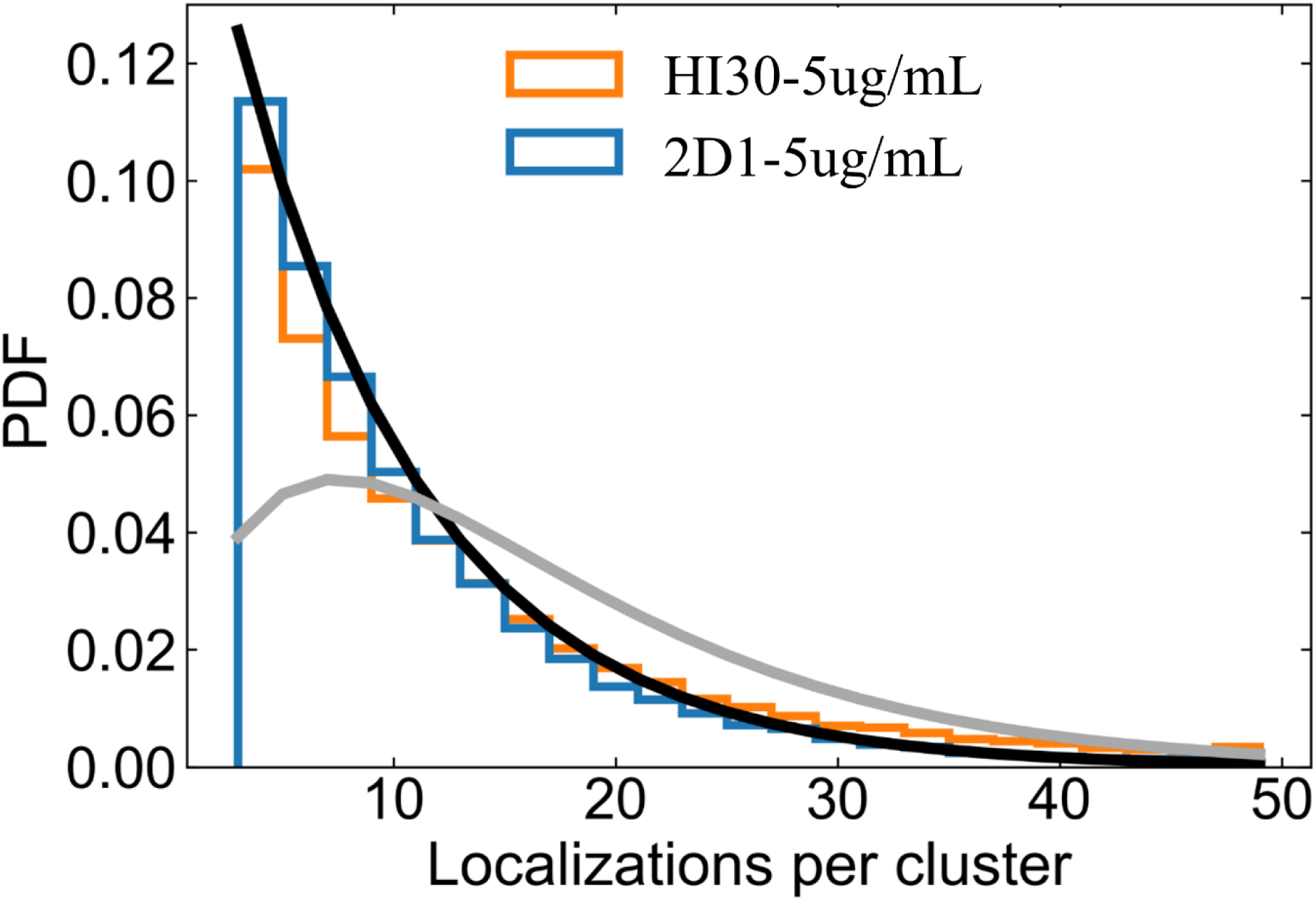
Probability distribution for the number of localizations per cluster for *d*STORM data of Jurkat T cells of low passage number immunostained for CD45 with various antibodies. *d*STORM localization data was clustered with a DBSCAN algorithm. Histograms of the number of localizations per cluster for CD45-2D1 (blue) and CD45-HI30 (orange) are displayed as probability density function (PDF). For comparison, theoretical expectations are shown for a monomeric distribution (black) and a dimeric distribution (gray) assuming an average of 8 localizations per antibody. Both, CD45-2D1 and CD45-HI30 show a pure monomeric distribution with no indication for dimeric contributions. In contrast to the data in Fig. 1C,E and Fig. 2C these cells have passed fewer passages and thus are closer to a physiological state.

**Fig. S8.**
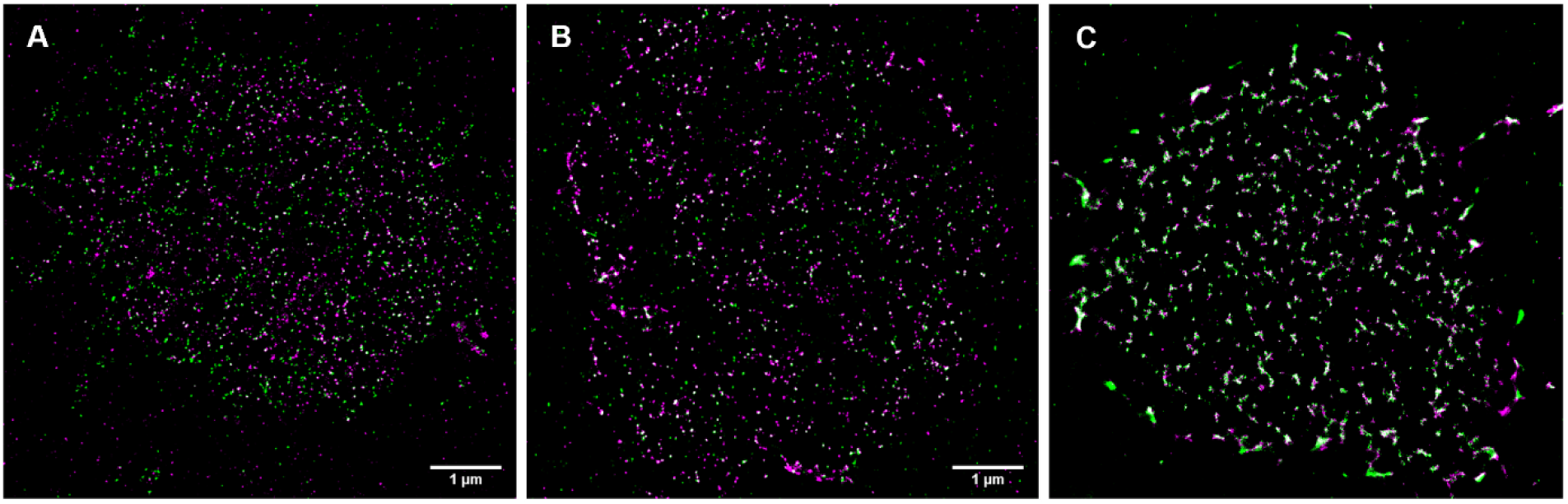
Artificial clustering is induced by additional incubation with a secondary antibody on live cell stained Jurkat T cells on ice. **A-C.** Cells were initially stained for 30 minutes with a mouse anti-CD45-AF647 (magenta) antibody, followed by an additional incubation with goat-anti-mouse (gam)-AF532 for 15 minutes (**A**), 30 minutes (**B**), and 1 hour (**C**) prior to fixation. While 15 min incubation resulted in only a few colocalization events with no significant clustering, 30 min incubation induces the appearance of nanoclusters that are even more pronounced after incubation for 1 h. Scale bars: 1 µm.

**Fig. S9.**
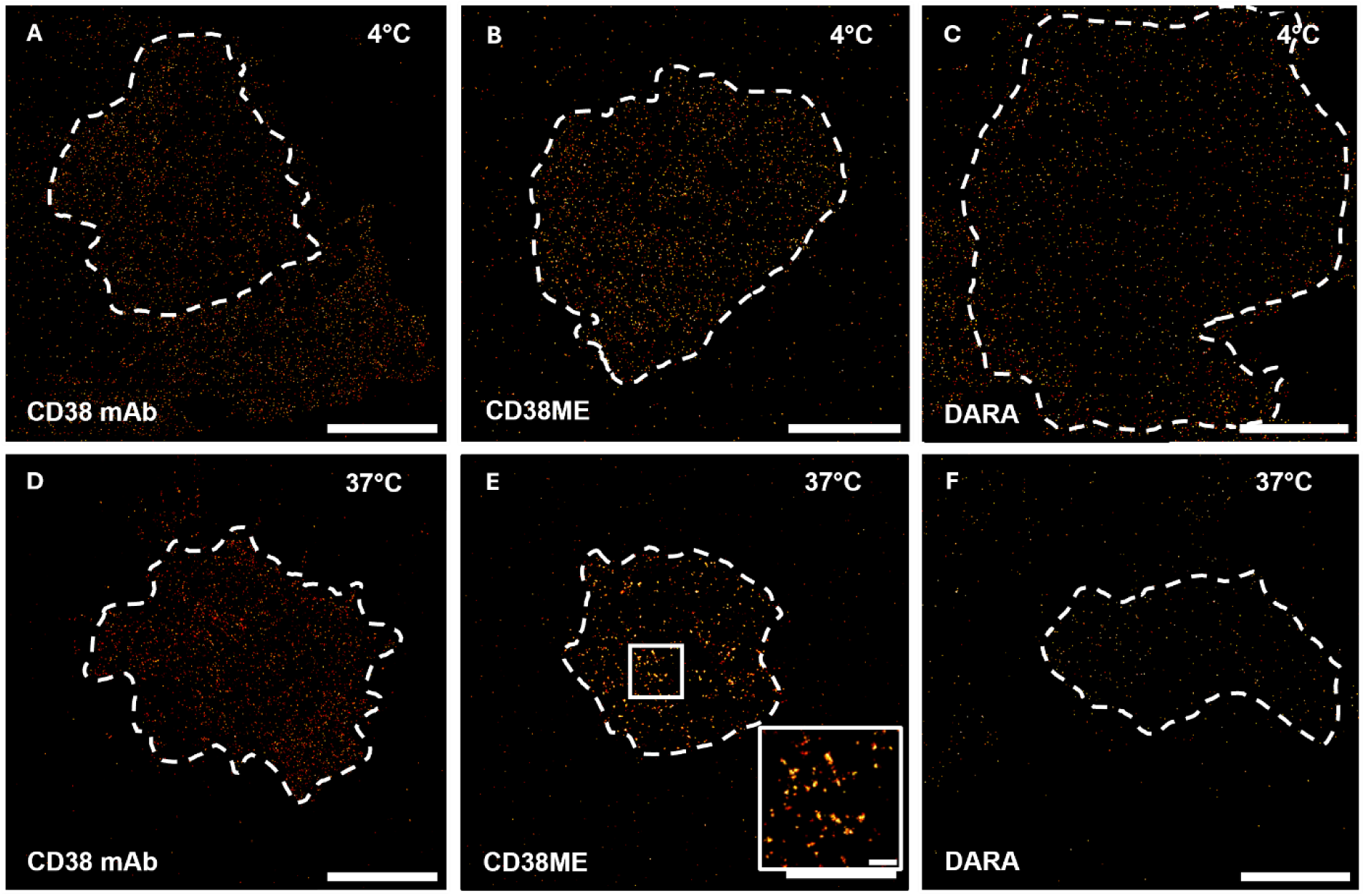
Influence of different monoclonal and polyclonal antibodies on receptor clustering. **A,D.** The monoclonal anti-CD38 antibody does not induce clustering of CD38 on OPM-2 cells at 4°C (**A**) and 37°C (**E**). **B,E.** Similarly, the polyclonal anti-CD38 antibody ME does not induce clustering at 4°C (**B**), but clustering becomes apparent when staining is performed at 37°C (**E**). **C,F.** The therapeutical monoclonal anti-CD38 antibody Daratumumab (DARA) shows no clustering of CD38 at 4°C (**C**) and 37°C (**F**). Scale bars, 2 µm, magnification 500 nm.

**Fig. S10.**
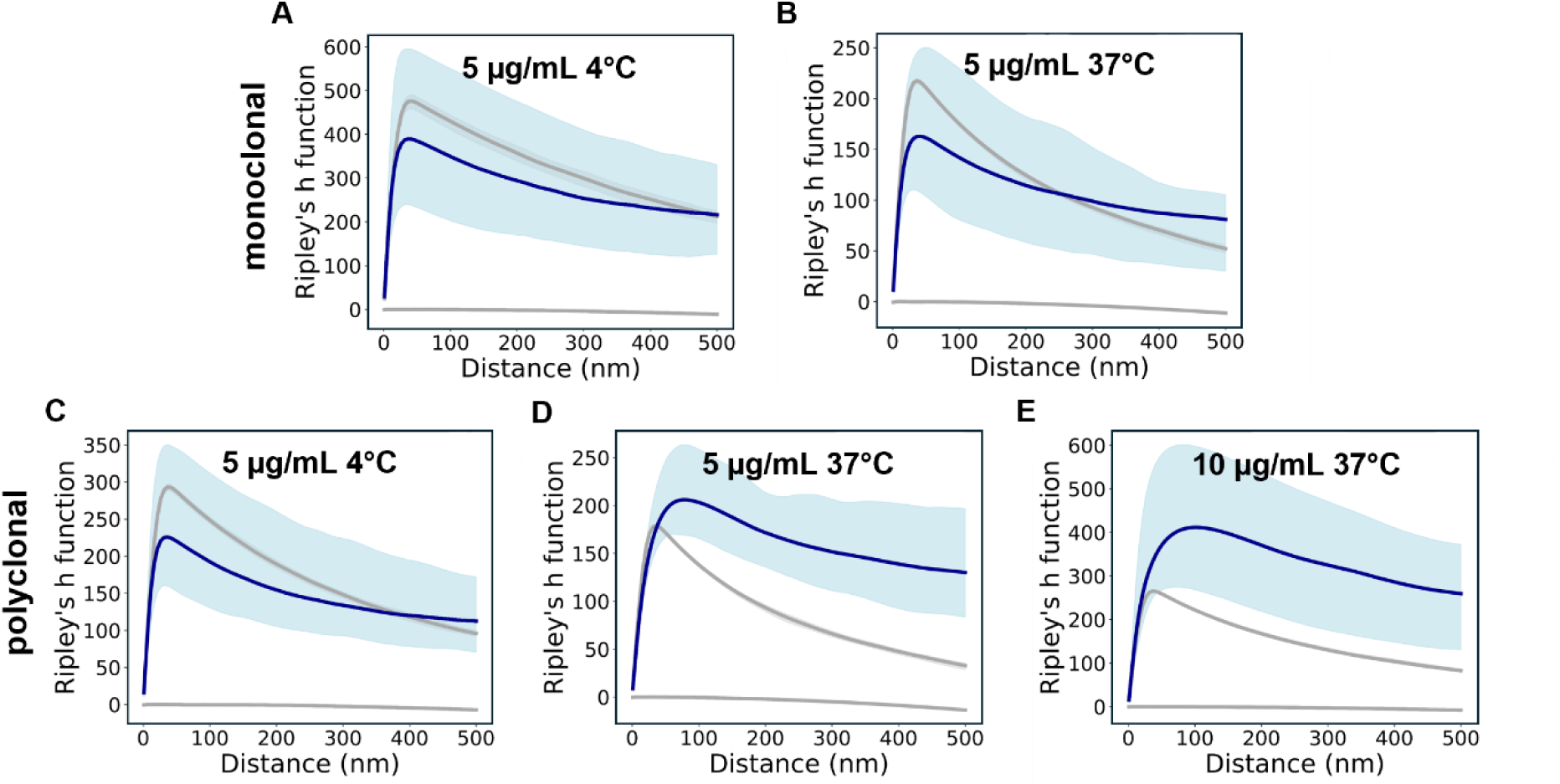
Ripley’s H-function analysis for *d*STORM data of OPM-2 cells stained for CD38. **A-C.** No clustering was detected for monoclonal antibody staining at either 4°C (**A**) or 37°C (**B**), nor with a polyclonal antibody staining at 4°C (**C**). **D-E.** Clear clustering was observed using a polyclonal antibody for staining at 37°C (**D**), which became even more pronounced at a concentration of 10 µg/mL (**E**). For comparison with experimental data from 6 to 10 cells (blue), Ripleýs H function for simulated data with spatial distributions following complete spatial randomness (lower grey lines) or a clustered Neyman-Scott process (upper grey lines) in identical ROIs are displayed with 95% confidence intervals (light blue and gray regions; if not shown confidence intervals are smaller than the line width).

**Fig. S11.**
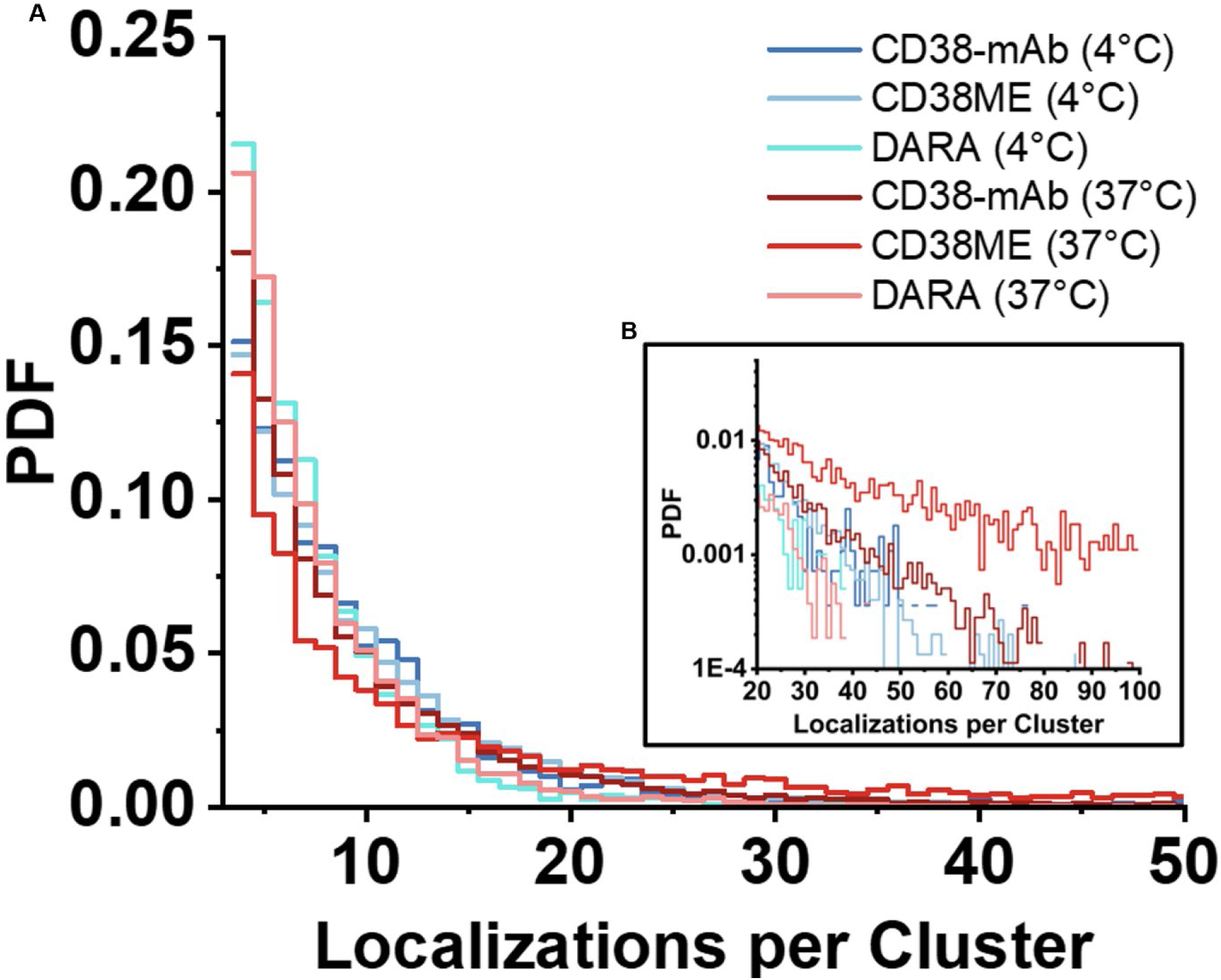
Probability distribution of OPM-2 cells stained for CD38. No clustering was detected with monoclonal antibody (mAb) or DARA staining at either 4°C or 37°C (**A**), nor with a polyclonal antibody staining at 4°C (CD38-ME). In contrast, clustering was observed with polyclonal antibody staining at 37°C (CD38-ME), showing not only reduced quantities of monomers but also an increase in higher order oligomers containing more localizations per cluster (**B**).

**Fig. S12.**
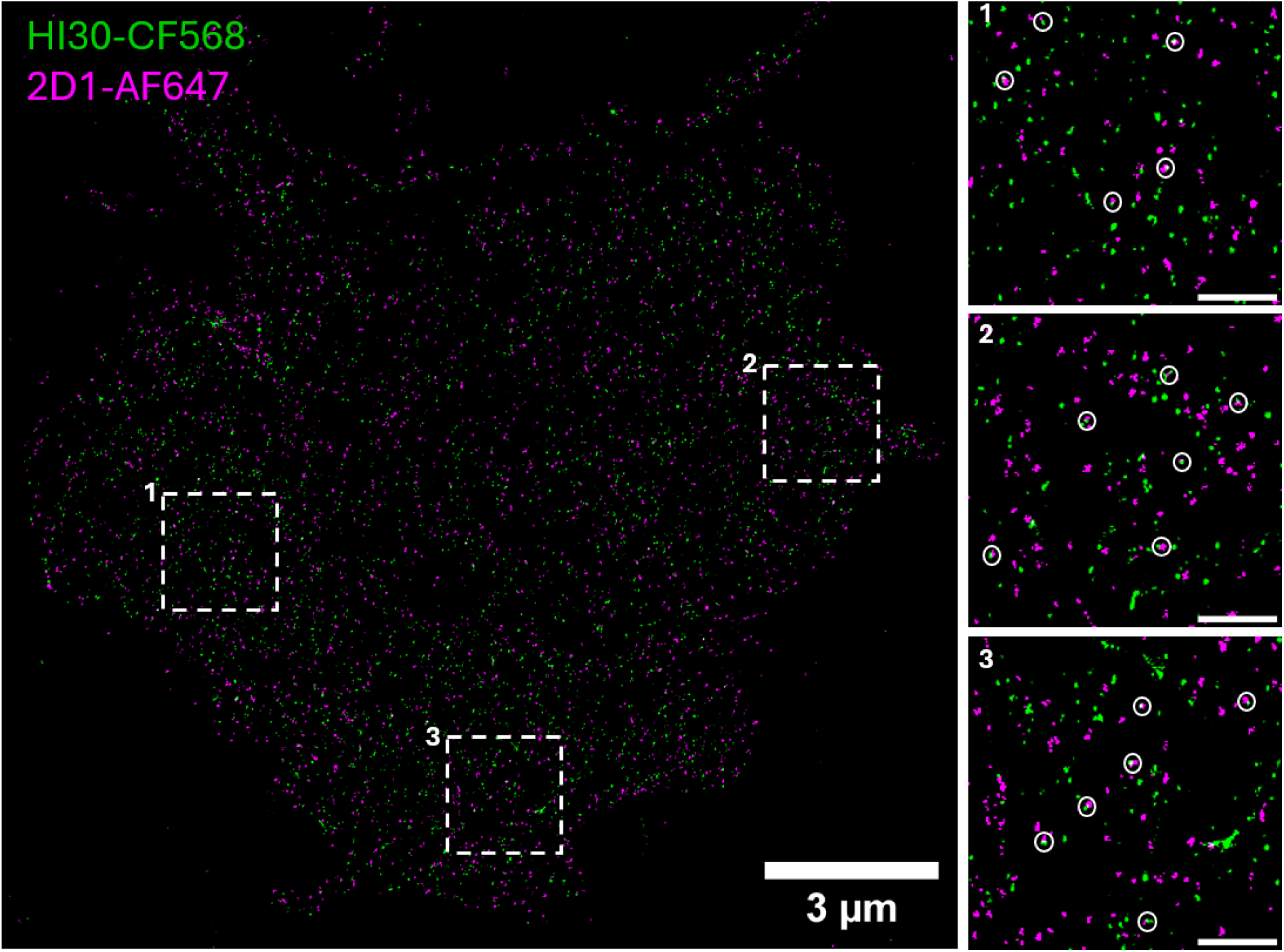
Two-color *d*STORM image of CD45-AF647 (clone 2D1, magenta) and CD45-CF568 (clone HI30, green). Jurkat T cells were stained following the live cell staining protocol with clone 2D1 for 15 minutes, followed by an additional incubation with clone HI30 for another 15 minutes prior to fixation. Several colocalizations are observed, representing either a single CD45 receptor stained with two antibodies or a potential homodimer (white circles). Scale bars of magnifications, 500 nm.

**Fig. S13.**
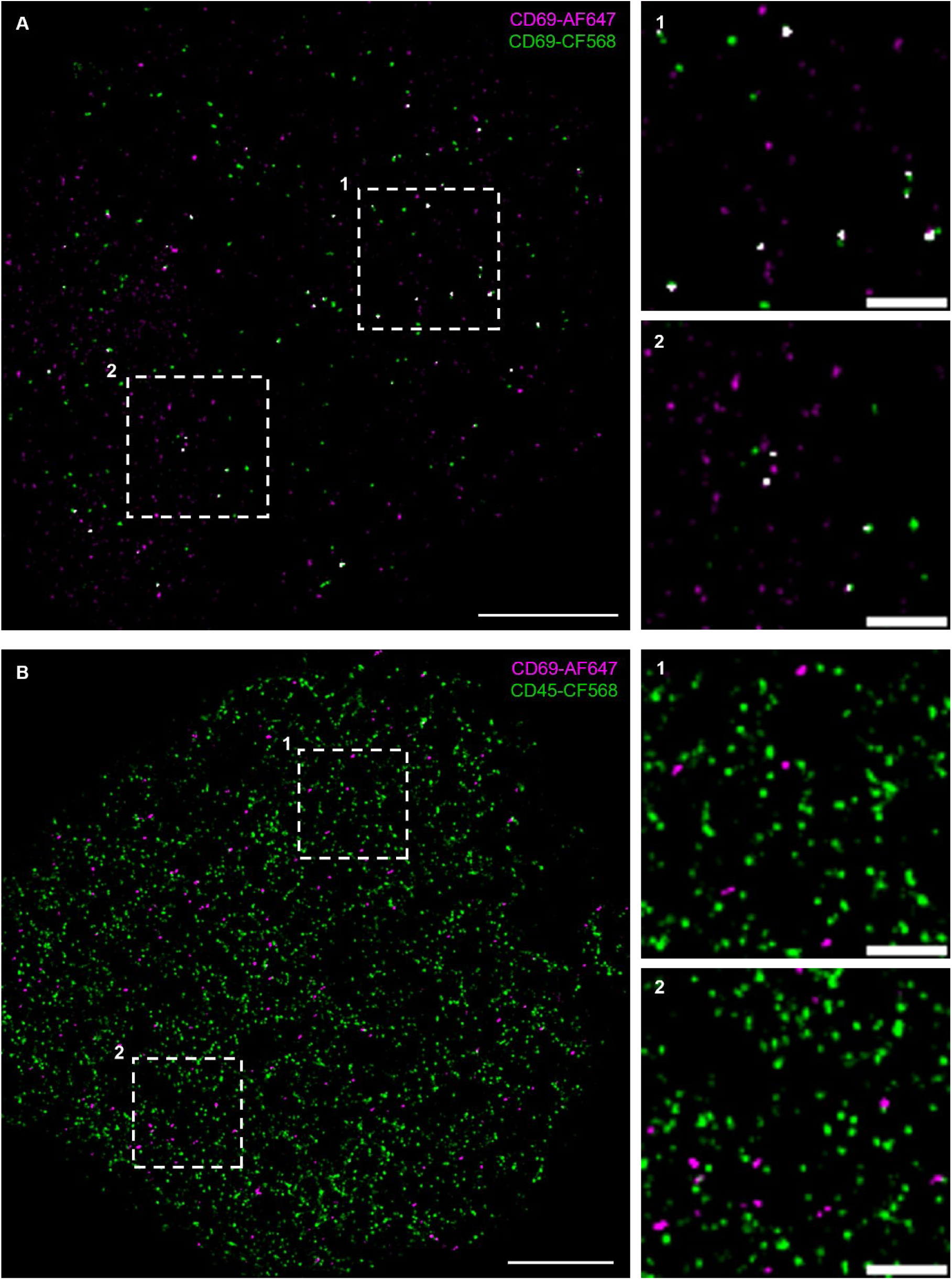
Two-color *d*STORM enables identification of possible homodimers. Jurkat T cells were stained following the live cell staining protocol with **A,** CD69 antibodies labeled with AF647 or CF568 simultaneously or **B,** CD69-AF647 and CD45-CF568 prior to fixation. Several colocalizations are observed for the homodimer CD69 (white), while CD69 and CD45 showed no colocalization at all. Scale bars: 2 µm, magnifications 500 nm.

**Fig. S14.**
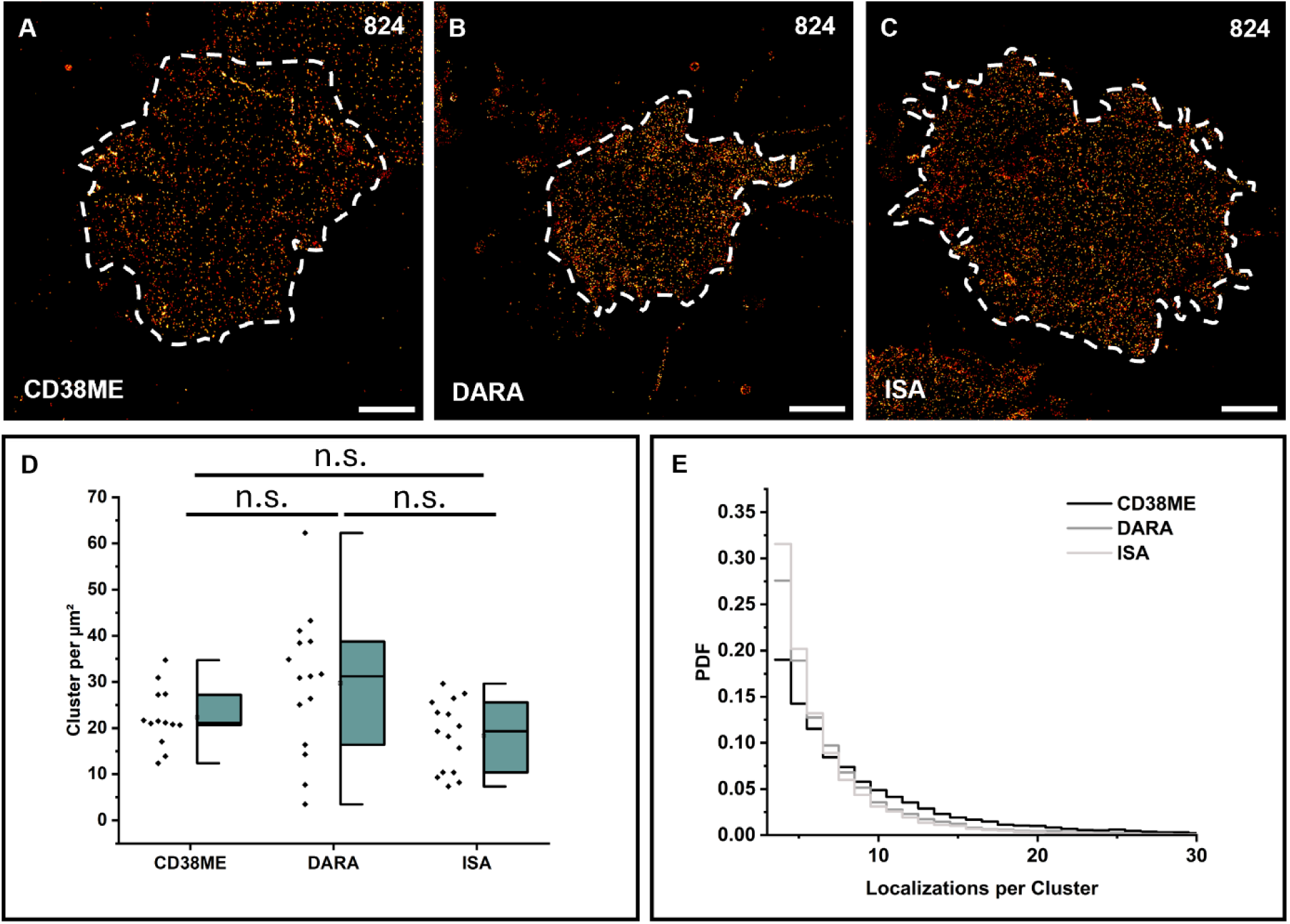
*d*STORM imaging and quantification of CD38 on tumor cells of a patient who became resistant to Daratumumab treatment. **A-C.** Representative *d*STORM images of the basal membrane of multiple myeloma cells stained with the polyclonal antibody CD38ME (**A**), Daratumumab (**B**) and Isatuximab (**C**) are shown for patient 824, who is responding to DARA treatment. **D.** CD38 localization clusters per µm^2^ detected for patient 824 using CD38ME, DARA and ISA labeled with AF647 (N=12-15 cells). **E.** Probability density functions (PDFs) of the number of localizations detected per cluster for the two different patients and antibodies. The abbreviation n.s. stands for not significant. Scale bars: 2 µm.

