## Supplementary Figures for "Single-molecule localization microscopy reveals the molecular organization of endogenous membrane receptors"

**The PDF file includes:**

**Figs. S1 to S14**

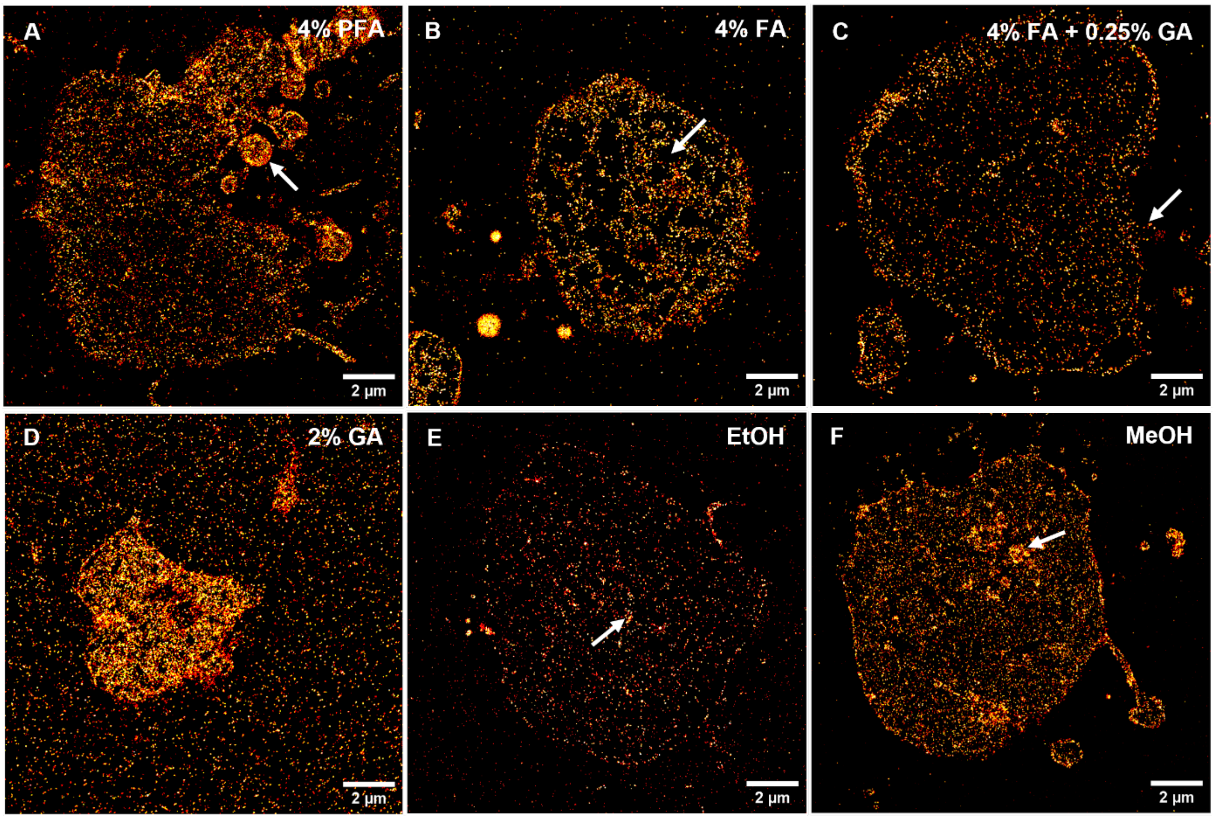


**Fig. S1. Representative *d*STORM images of the basal membrane of Jurkat T cells prefixed with various fixatives following staining with CD45-AF647 (HI30).** Fixation with 4% paraformaldehyde **A.** and 4% formaldehyde (**B**) leads to more pronounced apoptotic features, such as visible membrane blebbing or membrane disruptions (white arrows). **C.** A combination of 4% formaldehyde with 0.25% glutaraldehyde on the other side shows a high signal-to-noise ratio with only minor disruptions. **D.** Fixation with 2% glutaraldehyde induces high unspecific binding of the antibody to the glass surface. Cells fixed with 100% ethanol (**E**) and 100% methanol (**F**) exhibit a high signal-to-noise ratio, however, ethanol-fixed cells show reduced binding and intracellular staining. Arrows indicate the described features in each respective image.

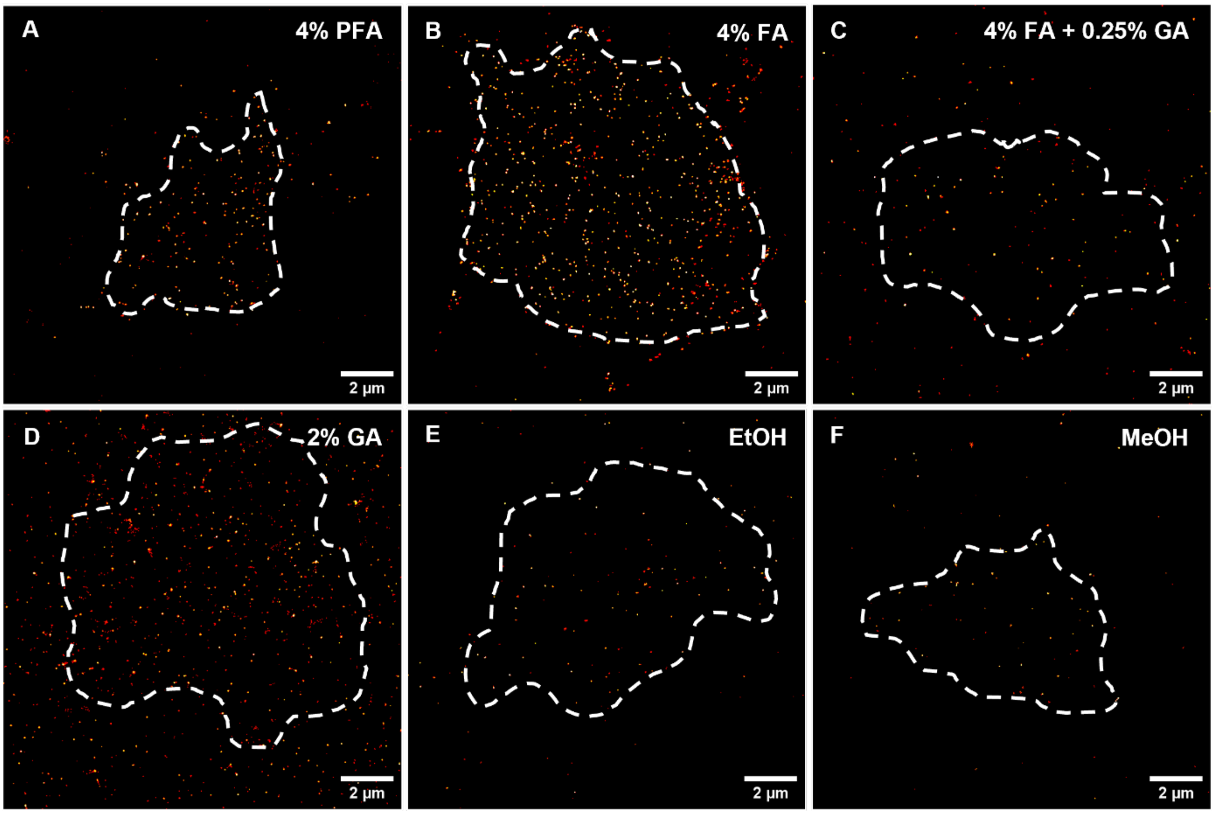


**Fig. S2. Representative *d*STORM images of the basal membrane of Jurkat T cells prefixed with various fixatives following staining with CD69-AF647.** Clear detection of CD69 molecules was observed with 4% paraformaldehyde (**A**) and 4% formaldehyde (**B**). However, protocols using 4% formaldehyde + 0.25% glutaraldehyde (**C**), 100% ethanol (**E**), and 100% methanol **(F)** resulted in failed or minor detection of CD69 expression. Additionally, a significantly higher amount of nonspecifically bound CD69 antibodies to the glass surface was observed with 2% glutaraldehyde prefixation (**D**).

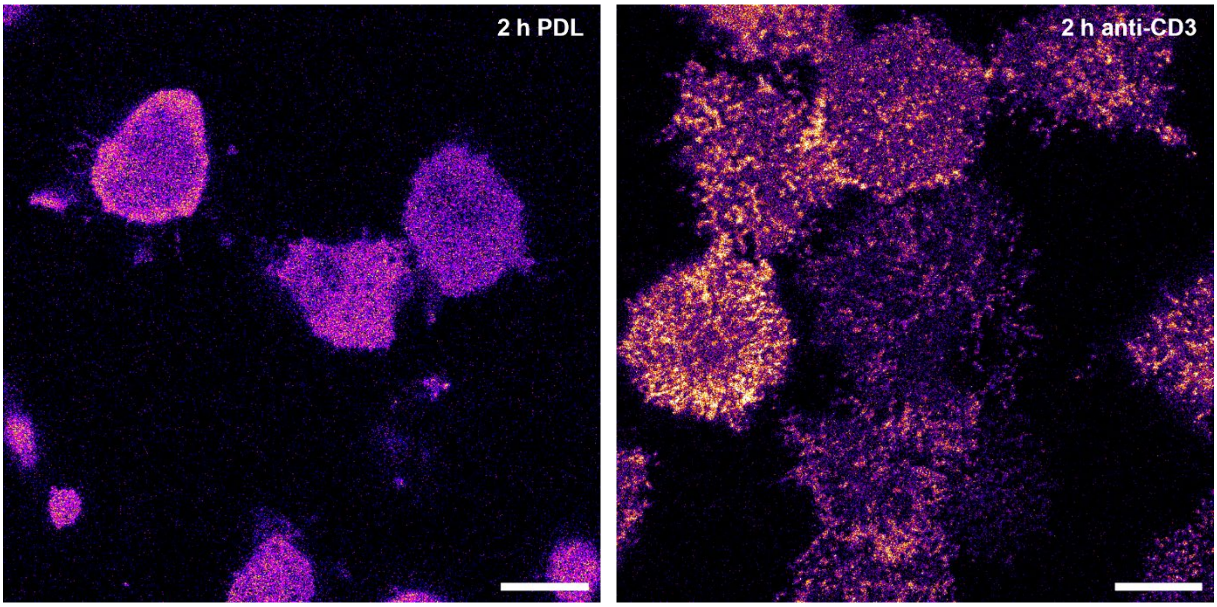


**Fig. S3. Confocal laser scanning images of Jurkat T ZAP70-GFP cells after 2 h on a PDL coated surface (left) and an anti-CD3-coated surface (right)**. Intracellular ZAP70-GFP remains homogeneously distributed on PDL, while binding to anti-CD3 antibodies activates the Jurkat T cells as indicated by ZAP70-GFP nanocluster formation.

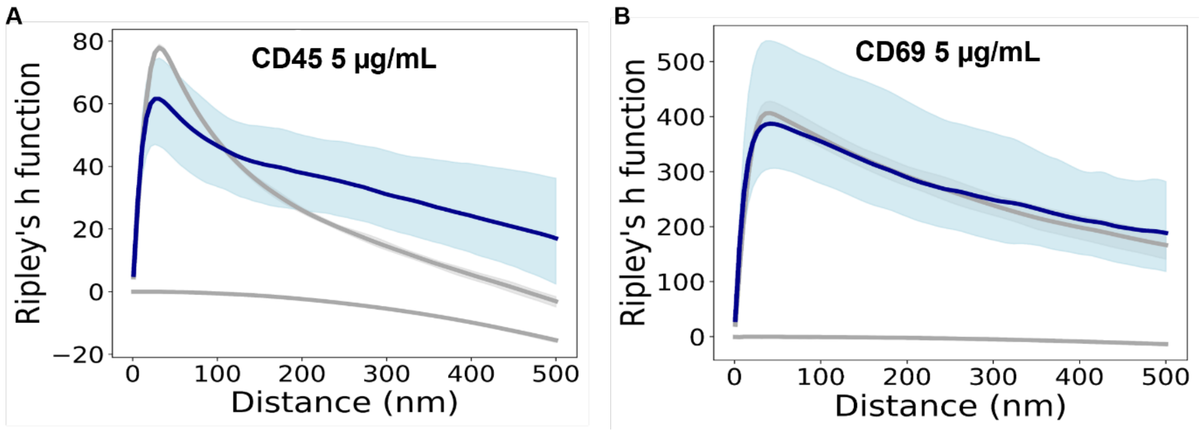


**Fig. S4. Ripley's H-function analysis for *d*STORM data of Jurkat T cells of low passage number immunostained for CD45 and CD69.** No significant clustering was observed for anti-CD45 (HI30) (**A**) and anti-CD69 (**B**) antibody staining on Jurkat T cells. Furthermore, no difference in cluster size between the two receptors was detected. For comparison with experimental data from 8 and 6 cells (blue), Ripley´s H function for simulated data with spatial distributions following complete spatial randomness (lower grey lines) or a clustered Neyman-Scott process (upper grey lines) in identical ROIs are displayed with 95% confidence intervals (light blue and gray regions; if not shown confidence intervals are smaller than the line width).

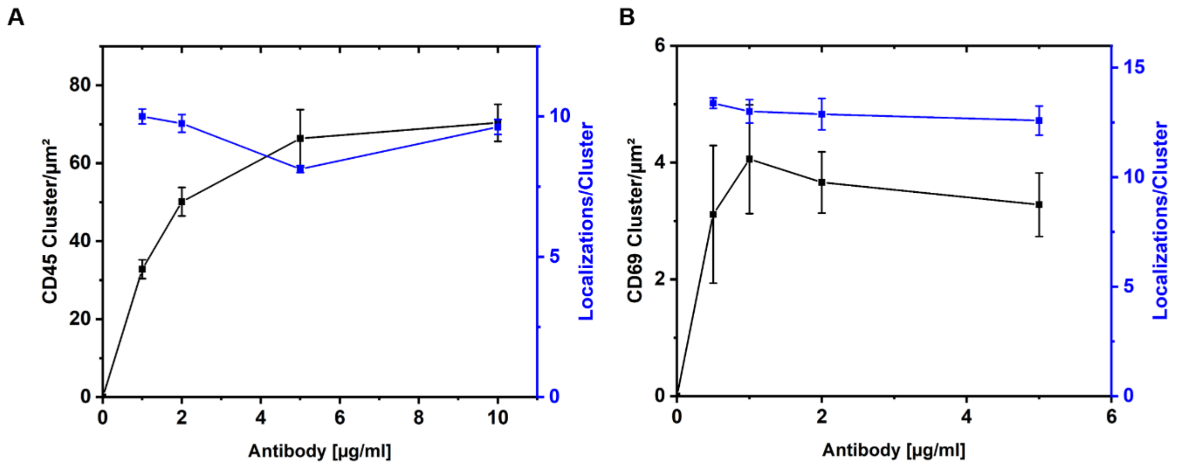


**Fig. S5. Average number of CD45 and CD69 localization clusters detected on the plasma membrane of Jurkat T cells at varying antibody concentrations. A,B,** The data indicate that an antibody concentration of 5 µg/mL and 2 µg/mL is sufficient to label all accessible CD45 (**A**) and CD69 (**B**) epitopes on the plasma membrane, respectively. Furthermore, the number of localizations detected per cluster remains consistent across varying concentrations of the antibodies, supporting the conclusion that the monoclonal CD45 (**A**) and CD69 (**B**) antibody does not induce artificial clustering.

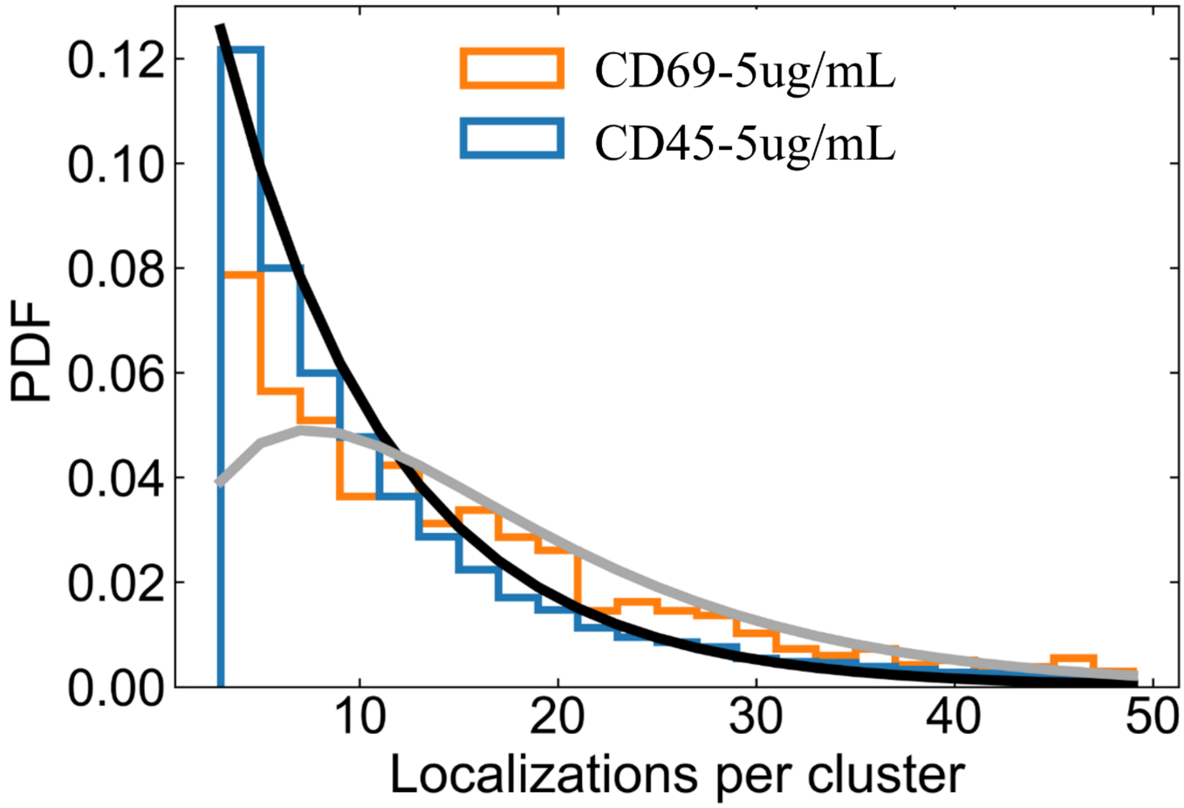


**Fig. S6. Probability distribution for the number of localizations per cluster for *d*STORM data of Jurkat T cells of low passage number immunostained for CD45 and CD69.** The same *d*STORM localization data as analyzed in fig. S4 was clustered with a DBSCAN algorithm. Histograms of the number of localizations per cluster for anti-CD45 (HI30) (blue) and anti-CD69 (orange) antibody staining on Jurkat T cells are displayed as probability density function (PDF). For comparison, theoretical expectations are shown for a monomeric distribution (black) and a dimeric distribution (gray) assuming an average of 8 localizations per antibody. For CD45 (HI30) the distribution resembles that of monomers. For CD69 a mixture distribution of monomers and a more pronounced dimeric population is observed.

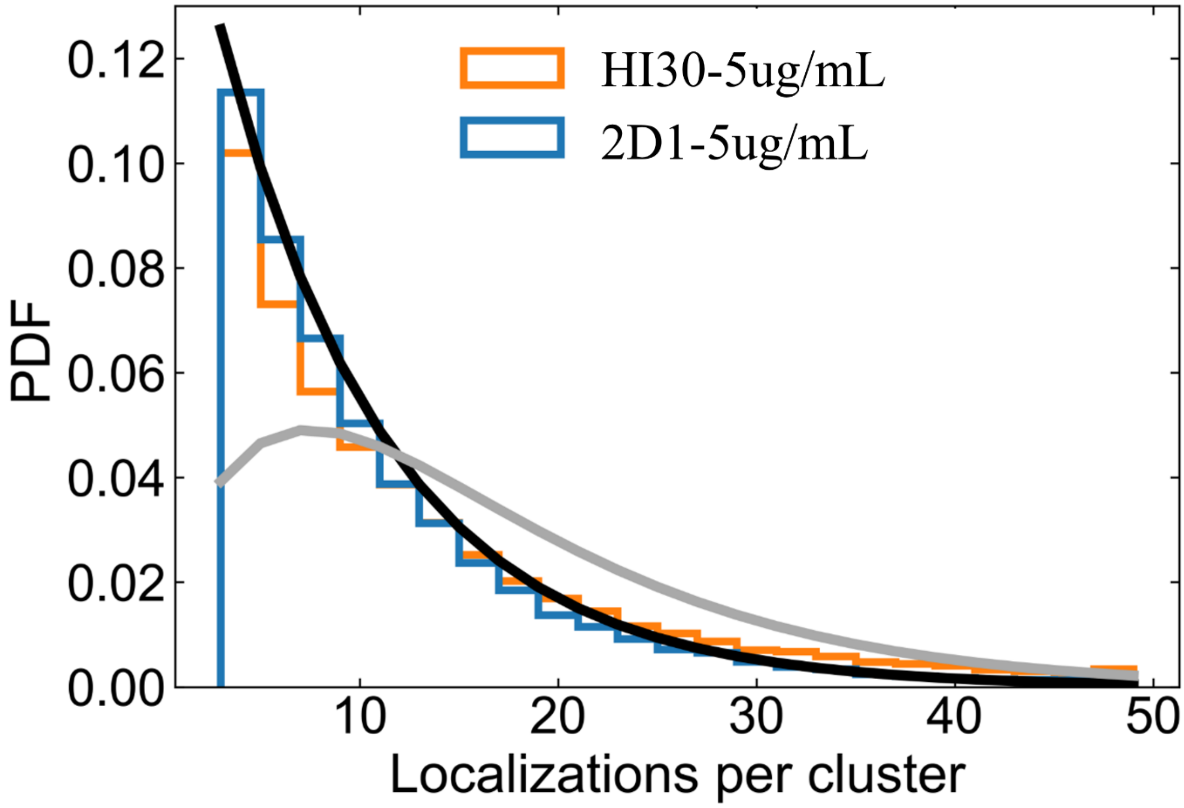


**Fig. S7. Probability distribution for the number of localizations per cluster for *d*STORM data of Jurkat T cells of low passage number immunostained for CD45 with various antibodies.** *d*STORM localization data was clustered with a DBSCAN algorithm. Histograms of the number of localizations per cluster for CD45-2D1 (blue) and CD45-HI30 (orange) are displayed as probability density function (PDF). For comparison, theoretical expectations are shown for a monomeric distribution (black) and a dimeric distribution (gray) assuming an average of 8 localizations per antibody. Both, CD45-2D1 and CD45-HI30 show a pure monomeric distribution with no indication for dimeric contributions. In contrast to the data in Fig. 1C,E and Fig. 2C these cells have passed fewer passages and thus are closer to a physiological state.

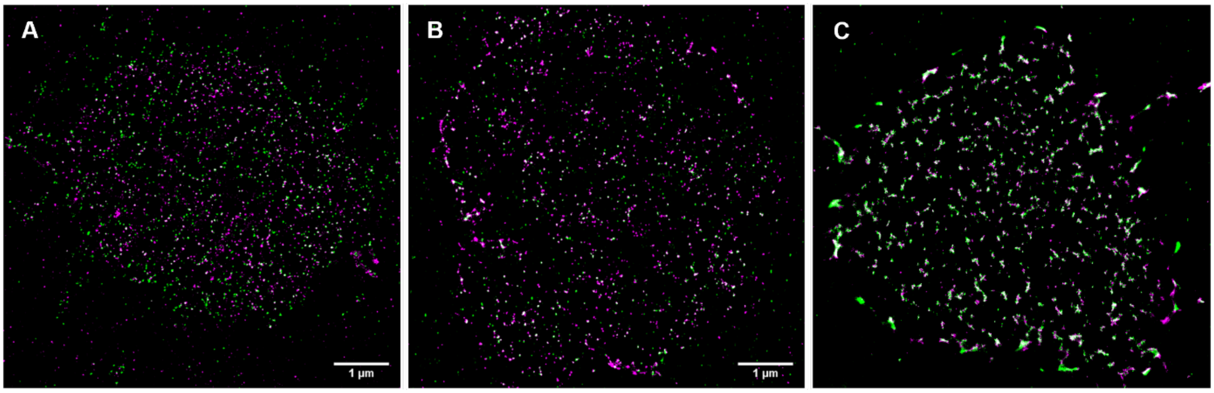


**Fig. S8. Artificial clustering is induced by additional incubation with a secondary antibody on live cell stained Jurkat T cells on ice. A-C.** Cells were initially stained for 30 minutes with a mouse anti-CD45-AF647 (magenta) antibody, followed by an additional incubation with goat-anti-mouse (gam)-AF532 for 15 minutes (**A**), 30 minutes (**B**), and 1 hour (**C**) prior to fixation. While 15 min incubation resulted in only a few colocalization events with no significant clustering, 30 min incubation induces the appearance of nanoclusters that are even more pronounced after incubation for 1 h. Scale bars: 1 µm.

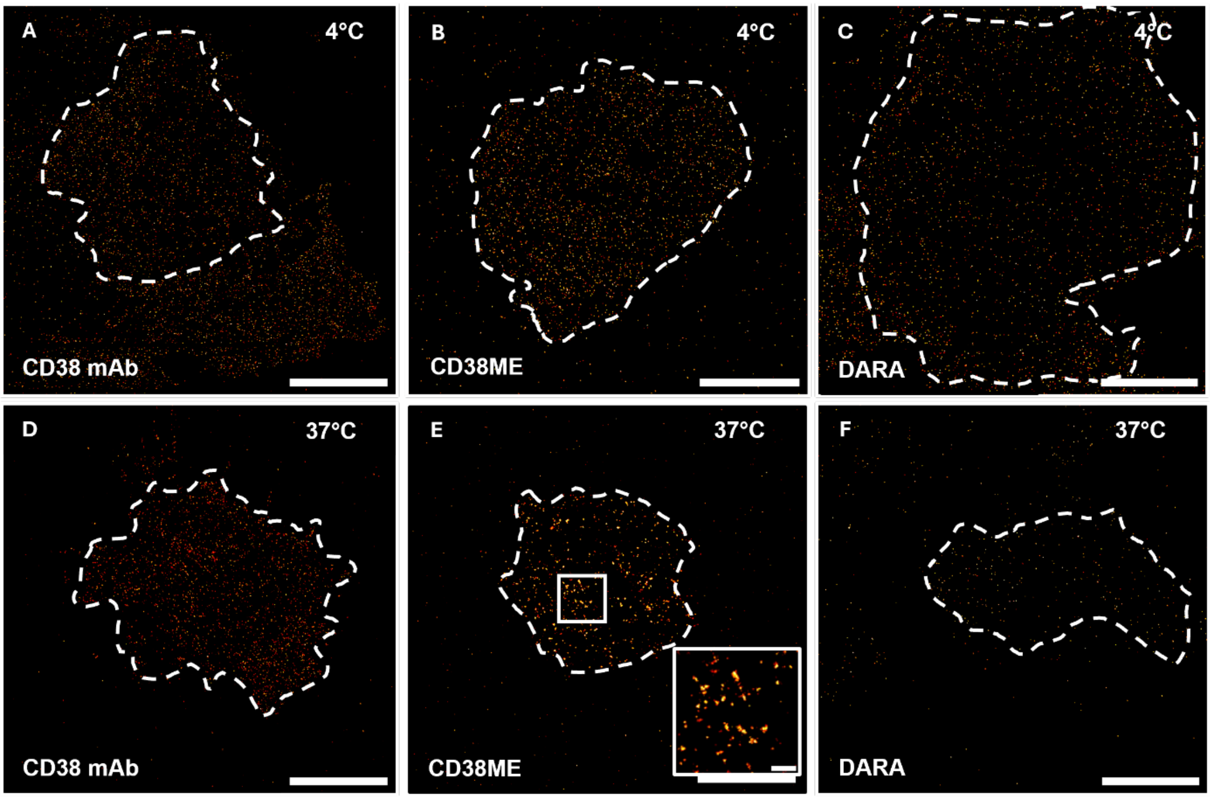


**Fig. S9. Influence of different monoclonal and polyclonal antibodies on receptor clustering. A,D.** The monoclonal anti-CD38 antibody does not induce clustering of CD38 on OPM-2 cells at 4°C (**A**) and 37°C (**E**). **B,E.** Similarly, the polyclonal anti-CD38 antibody ME does not induce clustering at 4°C (**B**), but clustering becomes apparent when staining is performed at 37°C (**E**). **C,F.** The therapeutical monoclonal anti-CD38 antibody Daratumumab (DARA) shows no clustering of CD38 at 4°C (**C**) and 37°C (**F**). Scale bars, 2 µm, magnification 500 nm.

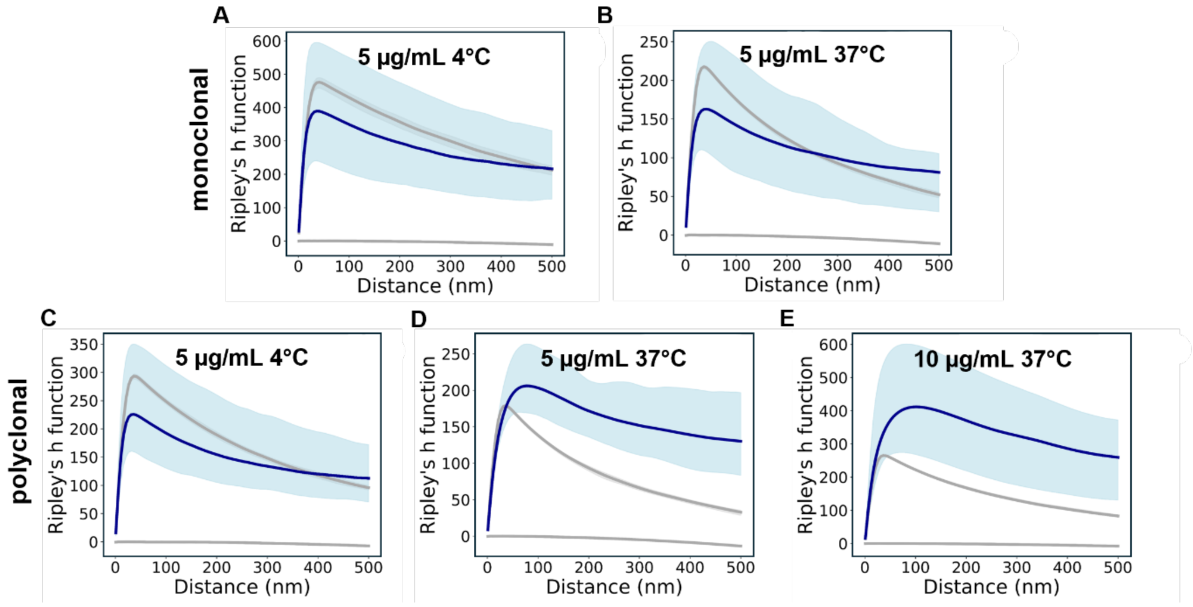


**Fig. S10 Ripley's H-function analysis for *d*STORM data of OPM-2 cells stained for CD38. A-C.** No clustering was detected for monoclonal antibody staining at either 4°C (**A**) or 37°C (**B**), nor with a polyclonal antibody staining at 4°C (**C**). **D-E.** Clear clustering was observed using a polyclonal antibody for staining at 37°C (**D**), which became even more pronounced at a concentration of 10 µg/mL (**E**). For comparison with experimental data from 6 to 10 cells (blue), Ripley´s H function for simulated data with spatial distributions following complete spatial randomness (lower grey lines) or a clustered Neyman-Scott process (upper grey lines) in identical ROIs are displayed with 95% confidence intervals (light blue and gray regions; if not shown confidence intervals are smaller than the line width).

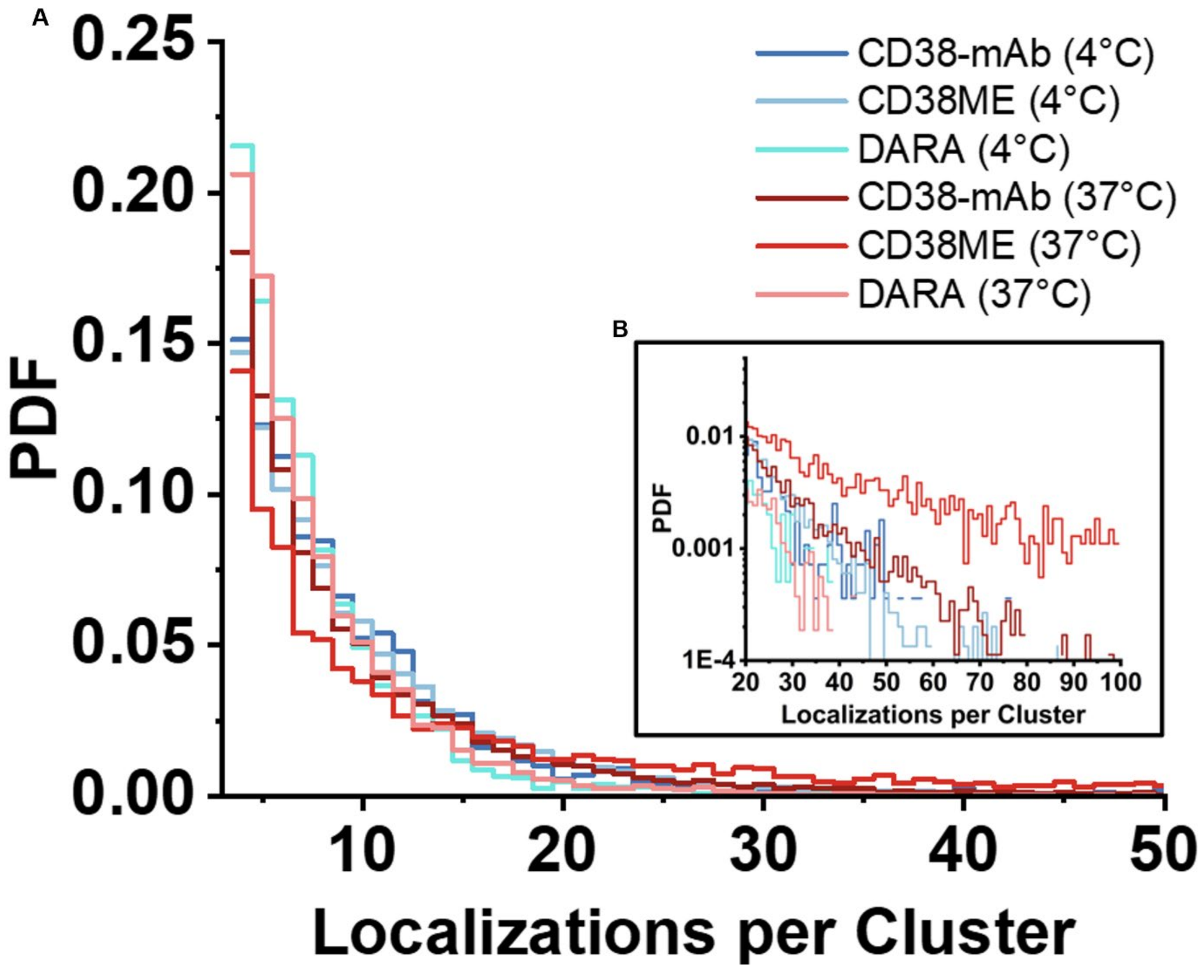


**Fig. S11. Probability distribution of OPM-2 cells stained for CD38.** No clustering was detected with monoclonal antibody (mAb) or DARA staining at either 4°C or 37°C (**A**), nor with a polyclonal antibody staining at 4°C (CD38-ME). In contrast, clustering was observed with polyclonal antibody staining at 37°C (CD38-ME), showing not only reduced quantities of monomers but also an increase in higher order oligomers containing more localizations per cluster (**B**).

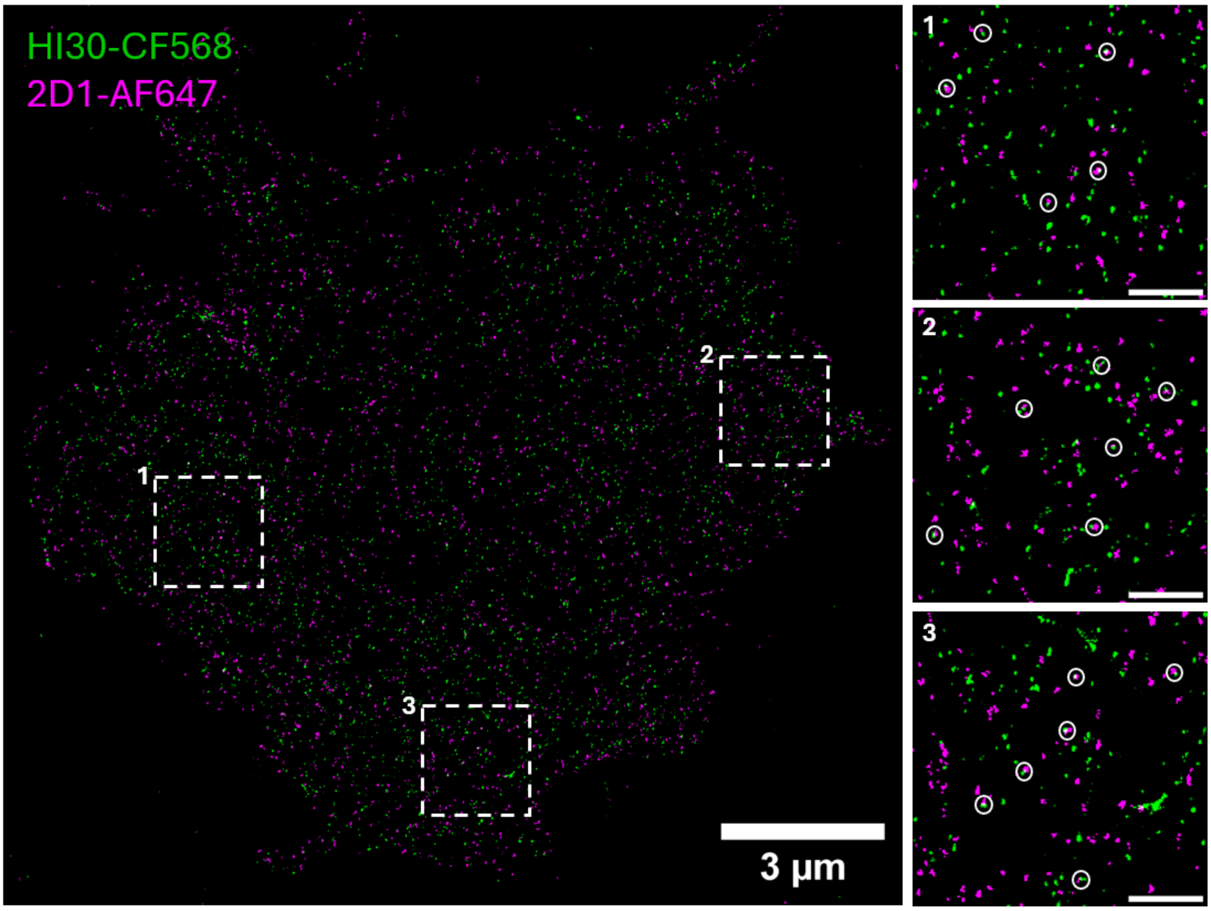


**Fig. S12. Two-color *d*STORM image of CD45-AF647 (clone 2D1, magenta) and CD45-CF568 (clone HI30, green).** Jurkat T cells were stained following the live cell staining protocol with clone 2D1 for 15 minutes, followed by an additional incubation with clone HI30 for another 15 minutes prior to fixation. Several colocalizations are observed, representing either a single CD45 receptor stained with two antibodies or a potential homodimer (white circles). Scale bars of magnifications, 500 nm.

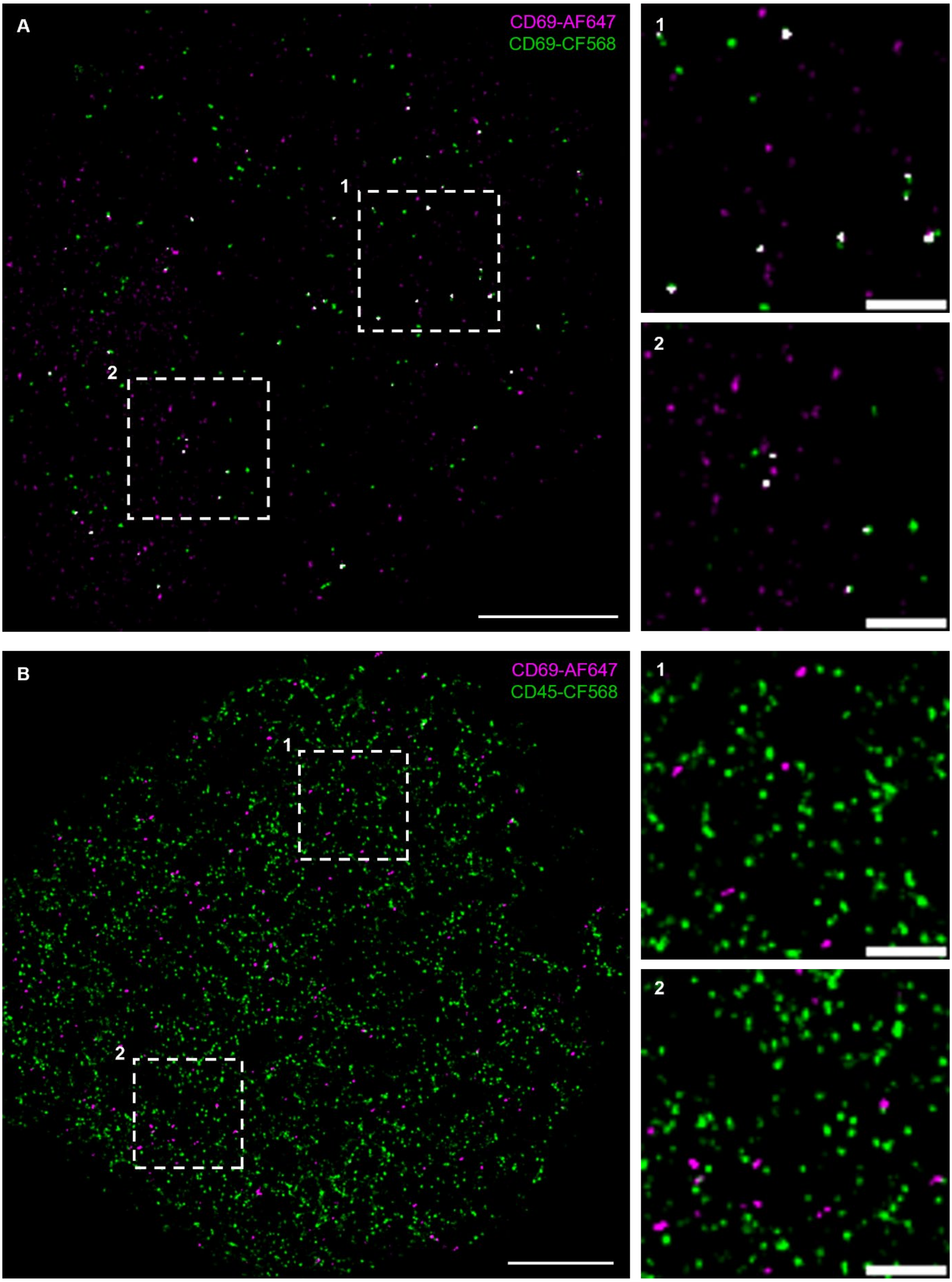


**Fig. S13. Two-color *d*STORM enables identification of possible homodimers.** Jurkat T cells were stained following the live cell staining protocol with **A,** CD69 antibodies labeled with AF647 or CF568 simultaneously or **B,** CD69-AF647 and CD45-CF568 prior to fixation. Several colocalizations are observed for the homodimer CD69 (white), while CD69 and CD45 showed no colocalization at all. Scale bars: 2 µm, magnifications 500 nm.

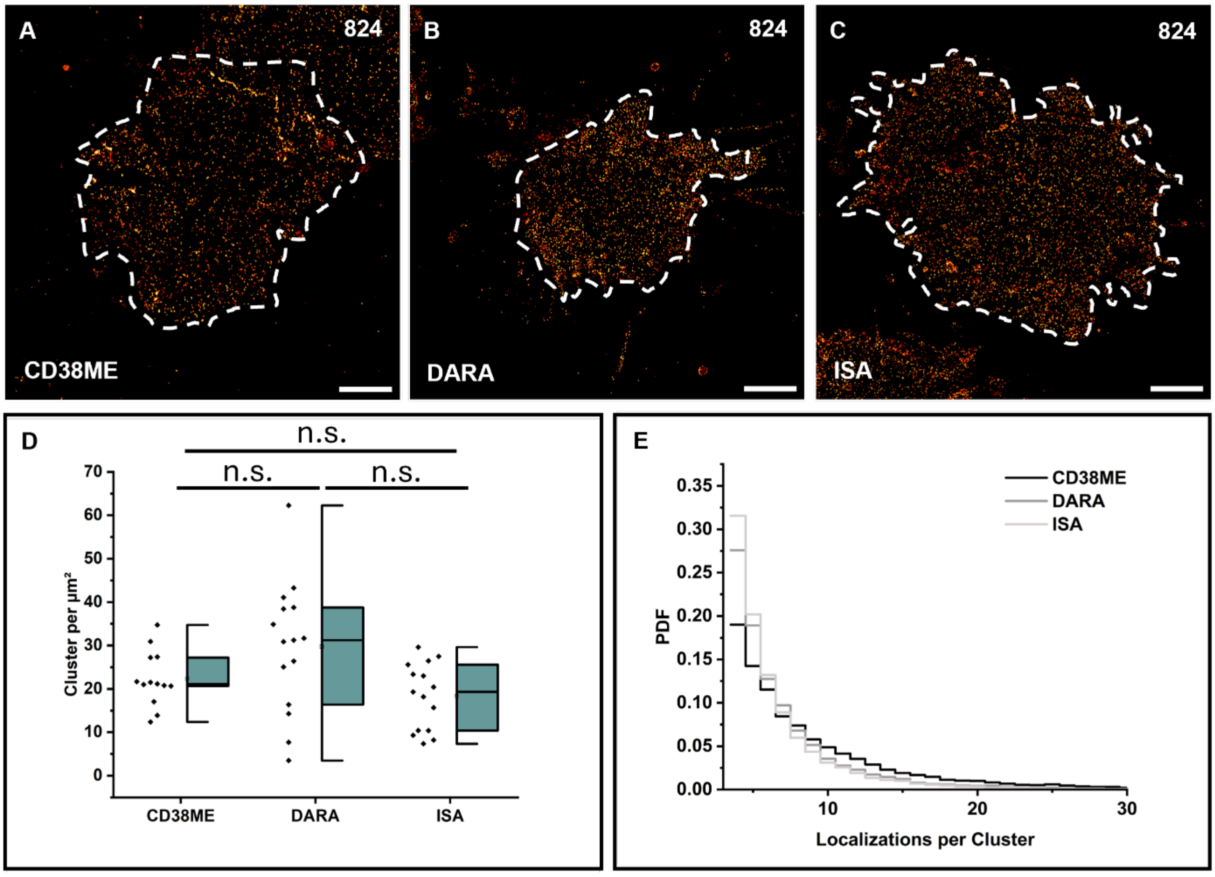


**Fig. S14. *d*STORM imaging and quantification of CD38 on tumor cells of a patient who became resistant to Daratumumab treatment. A-C.** Representative *d*STORM images of the basal membrane of multiple myeloma cells stained with the polyclonal antibody CD38ME (**A**), Daratumumab (**B**) and Isatuximab (**C**) are shown for patient 824, who is responding to DARA treatment. **D.** CD38 localization clusters per µm$^2$ detected for patient 824 using CD38ME, DARA and ISA labeled with AF647 (N=12-15 cells). **E.** Probability density functions (PDFs) of the number of localizations detected per cluster for the two different patients and antibodies. The abbreviation n.s. stands for not significant. Scale bars: 2 µm.
